# Distinct Co-Methylation Patterns in African and European Populations and Their Genetic Associations

**DOI:** 10.1101/2025.03.16.643544

**Authors:** Zheng (Joe) Dong, Nicole Gladish, Maggie P.Y. Fu, Samantha L. Schaffner, Keegan Korthauer, Michael S. Kobor

## Abstract

Human populations have substantial genetic diversity, but the extent of epigenetic diversity remains unclear, as population-specific DNA methylation (DNAm) has only been studied for ∼3.0% of CpGs. This study quantifies DNAm using whole-genome bisulfite sequencing (WGBS) and analyzes it alongside whole-genome genotype data to reveal a comprehensive picture of population-specific DNAm. Using a “co-methylated region” (CMR) approach, 36,657 CMRs were identified in 62 lymphoblastoid B cell line (LCL) WGBS samples, with validation in array data sets from 326 LCL samples. Between individuals of European and African ancestry, 101 CMRs exhibited population-specific DNAm patterns (Pop-CMRs), including 91 Pop-CMRs not found in previous investigations, which spanned genes (e.g., *CCDC42*, *GYPE*, *MAP3K20*, and *OBI1*) related to diseases (e.g., malaria infection and diabetes) with different prevalence and incidence rates between populations. Over half of the Pop-CMRs were asscoated with genetic variants, displaying population-specific allele frequencies and primarily mapping to genes involved in metabolic and infectious processes. Additionally, subsets of Pop-CMRs could be applicable in East Asian populations and peripheral blood-based tissues. This study provides insights into DNAm differences across the genome between populations and explores their associations with genetic variants and biological relevance, advancing our understanding of epigenetic roles in population specificity.

## Introduction

Human populations defined on the basis of ancestry have extensive phenotypic and molecular diversity as a consequence of environmental adaptations, including differences in immune function and disease susceptibility [1,2]. In the context of evolutionary biology, genetic variation is important in the adaptation of individual human populations and contributes to between-population phenotypic differences [2–5]. Beyond genetics, epigenetic modifications may contribute to variations in integral biological processes that are not fully explained by genetic variation, as epigenetic variation has been shown to play roles in development, disease, and responses to numerous environmental conditions [6–8]. DNA methylation (DNAm) is a highly stable and widespread epigenetic mark that has been widely studied in the context of inter-population epigenetic differences and the population-specific DNAm landscape [9–17]. For example, in lymphoblastoid B cell line (LCL) samples from five diverse human populations, population-specific DNAm derived from measurement of 485,000 CpGs mirrors genetic diversity, and can better reflect genetic influence than measurements at the transcriptomic level [11].

However, these previous studies can be improved upon with more advanced technologies and analytical approaches. First, all were investigated using various iterations of sparse DNAm arrays that cover only a small fraction of the roughly 28,000,000 CpG sites in the human genome. For example, the majority of these previous epigenome-wide association studies (EWASs) for human population specificity were conducted using the HM450K array, which covers ∼1.7% of genomic CpGs, thus limiting the assessment of genome-wide DNAm changes between populations and exploration of the genetic impact on the population specificity of DNAm. In contrast to DNAm arrays with relatively sparse CpG coverage, whole-genome bisulfite sequencing (WGBS) is a powerful method of assessing DNAm patterns throughout the full human genome as it can cover up to ∼95% of all genomic CpG sites (Li and Tollefsbol 2021). Increased coverage provides a more accurate representation of the methylation state and captures more nuance to any existing population-specific variation. Second, these studies focused on the population specificity of individual CpG sites. However, the deposition, maintenance, and removal of DNAm are predominantly controlled by DNA methyltransferases and ten-eleven translocation (TET) methylcytosine dioxygenases, which catalyze methylation alterations regionally rather than at the single-site level, forming regions of CpGs with correlated methylation status [18,19]. As genomic regions containing regulatory elements, such as gene promoters and CpG islands, are characterized by high CpG density, region-based analyses is likely of greater functional utility than single-CpG-based studies [20,21]. Region-based analyses can also alleviate the additional testing burden that attributes increasing, by 30-fold, the number of CpGs measured via data reduction by combining numerous adjacent CpGs into single units to reduce the multiple testing space [21]. Our group has recently developed a method called Co-Methylation with genomic CpG Background (CoMeBack) which performs genome-wide identification of co-methylated regions (CMRs), defined as regions of adjacent CpG sites with highly correlated DNAm status across individuals [21]. In contrast to differentially methylated regions (DMRs) constructed using phenotypic information, CMR identification does not use any phenotypic information and instead depends only on the correlation structure of adjacent CpG sites [21,22]. Therefore, using a region-based approach to evaluate whole-genome DNAm differences measured using WGBS between human populations could improve the discovery for population-specific DNAm alterations and provide insight into biological processes involved with population diversity.

In this study, we quantified DNAm using WGBS data and analyzed these data using the CMR approach to obtain a comprehensive picture of population-specific DNAm. We identified unique population-specific DNAm patterns compared to previous array-based studies and observed a relation among population-specific DNAm, genetic variation, and potential biological relevance. This research broadened our understanding of the crucial roles of epigenetic variation in population specificity.

## Results

### Study population and design

To analyze DNAm at whole-genome levels, we utilized published data from 62 WGBS LCL samples, taken from 8 individuals of European (EUR) ancestry and 54 individuals of African (AFR) ancestry (data sources are described in Supplementary table S1). As a first step in our multi-layered approach, we identified and characterized CMRs across the whole-genome using these WGBS LCL samples, with validation in 326 HM450K samples and functional annotation and enrichment analysis (Fig. 1a, data sources are described in Supplementary table S2). We next discovered population-specific CMRs (Pop-CMRs) between EUR and AFR WGBS samples and compared them to previous array-based investigations. We also investigated their enrichments in biological pathways to reveal their potential functional relevance. Furthermore, the association between genetic variation and the population specificity of DNAm patterns and the role of genetic variants in Pop-CMRs were evaluated using matched genotype data from 54 AFR whole-genome sequencing (WGS) samples. Functional investigations of single nucleotide polymorphisms (SNPs) associated with Pop-CMRs were carried out by examining their overlap with genome-wide association study (GWAS) hits and enrichments in an array of biological pathways. We also explored the expansion of Pop-CMRs to decipher population specificity in East Asian (EAS) LCL samples (n = 3) and an independent blood-based WGBS data set (healthy leukocyte samples, EUR n = 17, AFR n = 6; healthy plasma samples, EUR n = 17, AFR n = 6) (data sources are described in Supplementary table S1). Such a multi-layered approach allowed us to provide insights into the whole-genome DNAm differences between human populations and their associations with genetic variants and biological processes.

**Figure 1.**
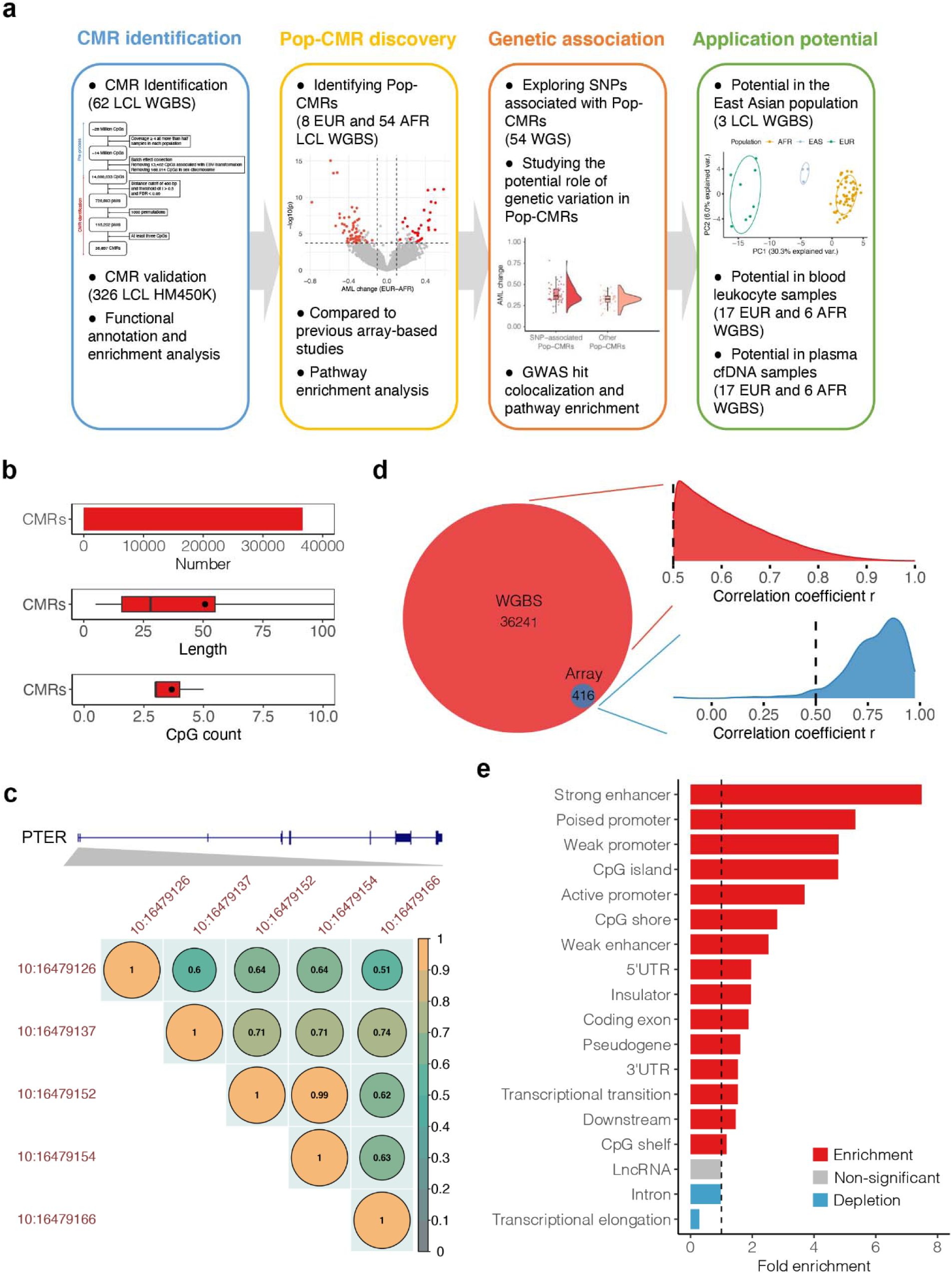
Identification and characterization of co-methylated regions (CMRs) in WGBS samples. **(a)** Flowchart of co-methylated region (CMR) generation and population specificity analysis in multiomics data. AFR, African ancestry; cfDNA, Cell-free DNA; EAS, East Asian ancestry; EUR, European ancestry; GWAS, genome-wide association study; LCL, lymphoblastoid B-cell line; Pop-CMR, population-specific CMR; WGS, whole-genome sequencing; WGBS, whole-genome bisulfite sequencing. **(b)** CMR characteristics, as evaluated by: total number of CMRs identified, distribution of CMR length, and distribution of CpG counts in the CMRs (top to bottom). **(c)** An example of a CMR containing five CpG sites located within the promoter of *PTER* (phosphotriesterase-related). The correlation values of adjacent CpG sites are colored as shown in the legend, where CpGs are included in a CMR if they were proximal and had *r* > 0.5 and FDR < 0.05. **(d)** Density plot of correlation coefficients amongst CpGs within a given CMR for CMRs identified in WGBS (EUR *n* = 8, AFR *n* = 54; top panel) and in a combined HM450K data set comprised of 326 LCL samples (EUR *n* = 157, AFR *n* = 169; bottom panel). **(e)** Significant enrichment (red) or depletion (blue) of CMRs in genomic elements (FDR < 0.05). A dashed vertical line indicates where fold enrichment equals 1. Fold enrichment was calculated based on random sampling repeated 1000 times. For each set of CMRs enriched in a certain genomic element, the same number and CpG density of regions were randomly selected from genomic sequences with a minimum mapping depth of 4 and without EBV transformation-associated and sex chromosome CpGs.

#### Co-methylated regions were extensive, reliable, and enriched in transcription regulatory elements

Data from 62 WGBS LCL samples, derived from 54 individuals of AFR ancestry (sampled from two different subpopulations—Yoruba in Nigeria (*n* = 8) and Gambian in Western Division – Mandinka (*n* = 46)) and 8 individuals of EUR ancestry (collected from two different countries—the USA (*n* = 5) and Sweden (*n* = 3)) were analyzed to identify CMRs (data sources are described in Supplementary table S1) [23–28]. The DNAm patterns of CpGs in sequencing reads were extracted from all samples and filtered for 4× coverage (Supplementary fig. S1). CpGs with DNAm changes associated with EBV immortalization of the LCLs were also excluded based on previous findings [28]. Subsequently, the local correlation structure of adjacent CpG methylation (*r*) was computed based on Pearson correlation analysis using all samples. Here, we defined CMRs as genomic regions containing at least three highly correlated CpG sites with CpG distance < 400 bp with a threshold of *r* > 0.5 and false discovery rate (FDR) < 0.05, which are stricter effect size and FDR threshold than used in previous studies to define CMRs (i.e., at least two correlated CpG sites with CpG distance < 400 bp and cutoff of *r* > 0.3 without the FDR threshold) [21,29]. To reduce the possible impacts of unbalanced sample sizes, we further validated the correlation structures of CpGs within each CMR using Pearson correlation by randomly selecting the same number of AFR samples as EUR samples with the same Pearson’s *r* and Benjamini–Hochberg FDR thresholds as above and repeating 1000 times (see Materials and Methods section “Identification of CMRs from LCL WGBS”).

36,657 CMRs were identified across the 62 WGBS samples, with an average block size of 50.81 bp with an average CpG count of 3.67 (Fig. 1b; the CMRs are listed in Supplementary table S3). The DNAm levels of the CpGs contained within a CMR are highly correlated by design as visualized in an example CMR (chr10:16479126–16479166) containing five adjacent CpGs located in the promoter of *PTER* (phosphotriesterase-related) ranging in *r* from 0.51 to 0.99 (Fig. 1c). Note that the majority of CMRs we identified were not covered by CpG probes in arrays (99.1% of the CMR were not represented on the HM450K array, which was mostly used in characterizing population-specific DNAm landscape), underscoring the importance of our study in enhancing our understanding of population-specific DNAm.

It remains to be seen how well CMRs can be reproduced using DNAm profiling techniques and DNA sources not represented in the data set used for CMR identification. To achieve this, we further validated these CMR correlation structures using a combined HM450K data set consisting of 326 LCL samples from three individual datasets (EUR *n* = 157, AFR *n* = 169 from GSE36369, GSE39672, and GSE40699; data sources are described in Supplementary table S2) [12,14,30]. With the same thresholds as above (*r* > 0.5 and FDR < 0.05), 94.8% of 416 CMRs that could be examined for correlation structures in the array (i.e., covering at least two CpGs; CpG *n* = 1033) were significantly correlated, suggesting the reliability of the CMRs identified in this study using WGBS samples (Fig. 1d).

Functional characterization of the genomic features encompassed within the identified CMRs was conducted by mapping them to 18 typical genomic elements retrieved from the UCSC Genome Browser and the GENCODE database [31]. The majority of CMRs (*n* = 29,792; 81.27%) were found to overlap with these genomic elements, especially in introns, strong enhancers, CpG shores, and weak enhancers (Supplementary table S4). Next, the enrichment of CMRs in each genomic element was calculated based on 1000 permutations compared to the same number of random regions with the same CpG count and similar CpG density (by restricting the maximum distance < 400 bp between two nearby CpGs as used in CMR identification). This found that CMRs exhibited significant enrichment in 15 genomic features, especially in typical regulatory elements, with > 5-fold enrichment in strong enhancers and poised promoters, whereas they exhibited significant depletion in regions of transcriptional elongation and introns (all FDR < 0.05) (Fig. 1e and Supplementary table S4). The significant enrichment in typical regulatory elements, such as promoters and enhancers, was consistent with findings from a previous study that identified CMRs using whole-blood HM450K array data [21]. This consistency underscored a major strength of the CMR approach, which might be more representative of functional relevance than individual CpG sites. Thus, by employing this approach, our study is likely to capture local coherent DNAm patterns that are either directly or indirectly associated with transcriptional regulation.

#### Comparable global CMR methylation profiles between EUR and AFR populations

To compare the global trends in DNAm landscapes between human populations, we analyzed genome-wide CMR methylation differences between EUR and AFR populations. DNAm levels for each CMR were calculated based on the average methylation level (AML), i.e., the average DNAm level of correlated CpG sites within the CMR. Globally, the two populations (EUR *n* = 8, AFR *n* = 54) had very similar DNAm patterns when assessing CMRs (cosine similarity = 0.98, average absolute value of DNAm difference = 8.87 × 10^−3^, Mann–Whitney U test *P* = 0.47). Principal component analysis (PCA) of CMR DNAm did not reveal global differences between the two populations (Supplementary fig. S2). Moreover, we did not observe any significant differences in DNAm between the two human populations across genomic features (cosine similarity ranged from 0.96 to 0.99, average absolute value of DNAm change ranged from 1.60 × 10^−4^ to 0.028 depending on genomic features, all Mann–Whitney U test FDR > 0.05) (Supplementary table S5). These results demonstrated a high degree of similarity in global DNAm patterns in CMRs between individuals of EUR and AFR ancestry, and suggested that there were minimal to no systematic technical artifacts, such as batch effects or variations in sample processing, affecting the comparison between samples from the two populations.

#### Identification of population-specific CMRs with a trend toward higher methylation in AFR compared to EUR

The main goal of this study was to test the hypothesis that subtle yet important differences in DNAm at the levels of CMRs existed between EUR and AFR populations. As such, we used linear regression adjusting for sex to identify population-specific CMRs (Pop-CMRs) (see Materials and Methods section “Detection and validation of Pop-CMRs”). We further tested that whether there were any systematic biases in our association results by estimating the inflation factor (λ) using the `BACON` R package, an established tool for estimation and correction of bias and inflation in epigenome- and transcriptome-wide association studies [32]. We observed an inflation factor of 1.08, which suggested that there was unlikely to be an influence of systemic bias. A strict AML difference threshold of 0.1 between EUR and AFR populations together with an FDR cutoff of 0.05 was applied to minimize the false positive rate [33]. As detailed in Materials and Methods section “Detection and validation of Pop-CMRs”, we further used the 1000 permutation test to evaluate and reduce the possible effects of unbalanced sample size and subpopulation structures, respectively. Thirty-five CMRs that passed the AML and FDR thresholds were removed due to the possible effects of the unbalanced sample size (*n* = 34) and subpopulations (*n* = 1).

Through this approach, 101 Pop-CMRs that displayed significant DNAm differences between the two populations were identified (Fig. 2a, 2b, and Table 1). Among them, 69 (68.3%) were more highly methylated in AFR samples compared to EUR samples (i.e., the methylation was higher in AFR by any magnitude) (Fig. 2c). This percentage was significantly higher than the 57.5% observed in AFR samples when randomly selecting from all CMRs (1000 permutation test *P* < 0.001) (Fig. 2c). This result was consistent with a previous array-based study, albeit with a smaller difference (10.8% in our study vs. 22.3% in the previous study) [9], potentially due to differences in cell types (LCL vs. monocytes) and/or DNAm profiling methods used (WGBS vs. EPIC). We next performed PCA of these 101 Pop-CMRs to obtain a comprehensive overview of the DNAm patterns across samples. The two populations were clearly distinct in the loading of the first principal component (PC), which accounted for 31.5% of the total variance (Fig. 2d). As expected, the first PC for Pop-CMRs showed better performance in population clustering and explained a 2.5-fold higher proportion of the total variance compared to random sets of CMRs on 1000 permutation analysis (31.5% vs. 12.5%, respectively, *P* < 0.001) (Fig. 2e). These findings suggested that the discovery of Pop-CMRs was likely driven by biological relevance rather than chance.

**Figure 2.**
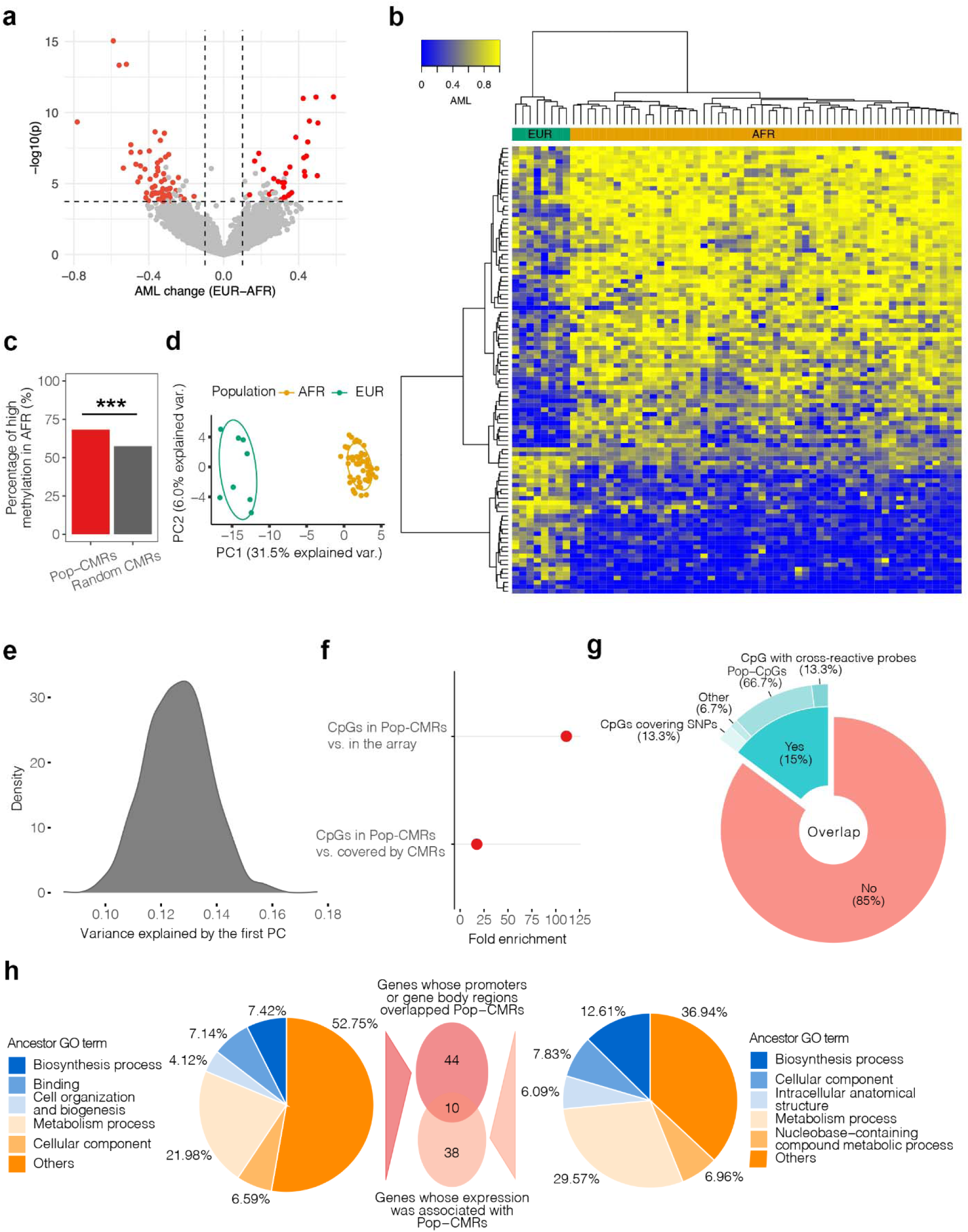
Identification and characterization of population-specific CMRs (Pop-CMRs) between European (EUR) and African (AFR) individuals. Data from all 62 WGBS samples (EUR *n* = 8, AFR *n* = 54) used in this study are shown. **(a)** Volcano plot of significant Pop-CMRs. In the volcano plot of Pop-CMRs, the signal detection results show the methylation differences (x-axis) and significance (y-axis) between EUR and AFR individuals adjusted for sex. The CMRs that are nonsignificant or affected by an unbalanced sample size and subpopulation structures are shown in gray. Each spot represents a CMR. **(b)** Unsupervised hierarchical clustering heat map using DNAm patterns from 101 Pop-CMRs. **(c)** Percentage of more highly methylated Pop-CMRs in AFR vs. EUR samples compared to random sets of CMRs. Random sampling was repeated 1000 times. *** *P* < 0.001. **(d)** Loadings using the first two PCs of 101 Pop-CMRs for each WGBS LCL sample on principal component analysis (PCA) color-coded by ancestry. **(e)** The density of proportion of variance was explained by the first PC from AMLs of randomly selected CMRs. The random sampling process was repeated 1000 times. Only the explained variances by first PC of random CMR sets are shown since it was calculated to compare the explained variances by first PC of Pop-CMRs. **(f)** Enrichment of CpGs in the array covered by Pop-CMRs showed statistically significant differences in DNAm between EUR and AFR populations compared to random CpG sets from CpGs in the array covered by CMRs and all CpGs in the array separately. The random sampling process was repeated 1000 times for each comparison. Both *P* < 0.001. **(g)** Pie-donut chart showing the overlap of 101 Pop-CMRs with array CpGs from HM27K, HM450K, and EPIC. Most of these overlapped CpGs were previously identified as population-specific (Pop-CpGs). **(h)** Pie charts showing Gene Ontology (GO) classification. Pathway enrichment analysis of 54 genes whose promoters or gene body regions overlapped Pop-CMRs is presented on the left. Pathway enrichment analysis of 48 genes whose expression was associated with Pop-CMRs in the iMETHYL database is presented on the right. The overlap is shown with the venn diagram in the center.

**Table 1.**
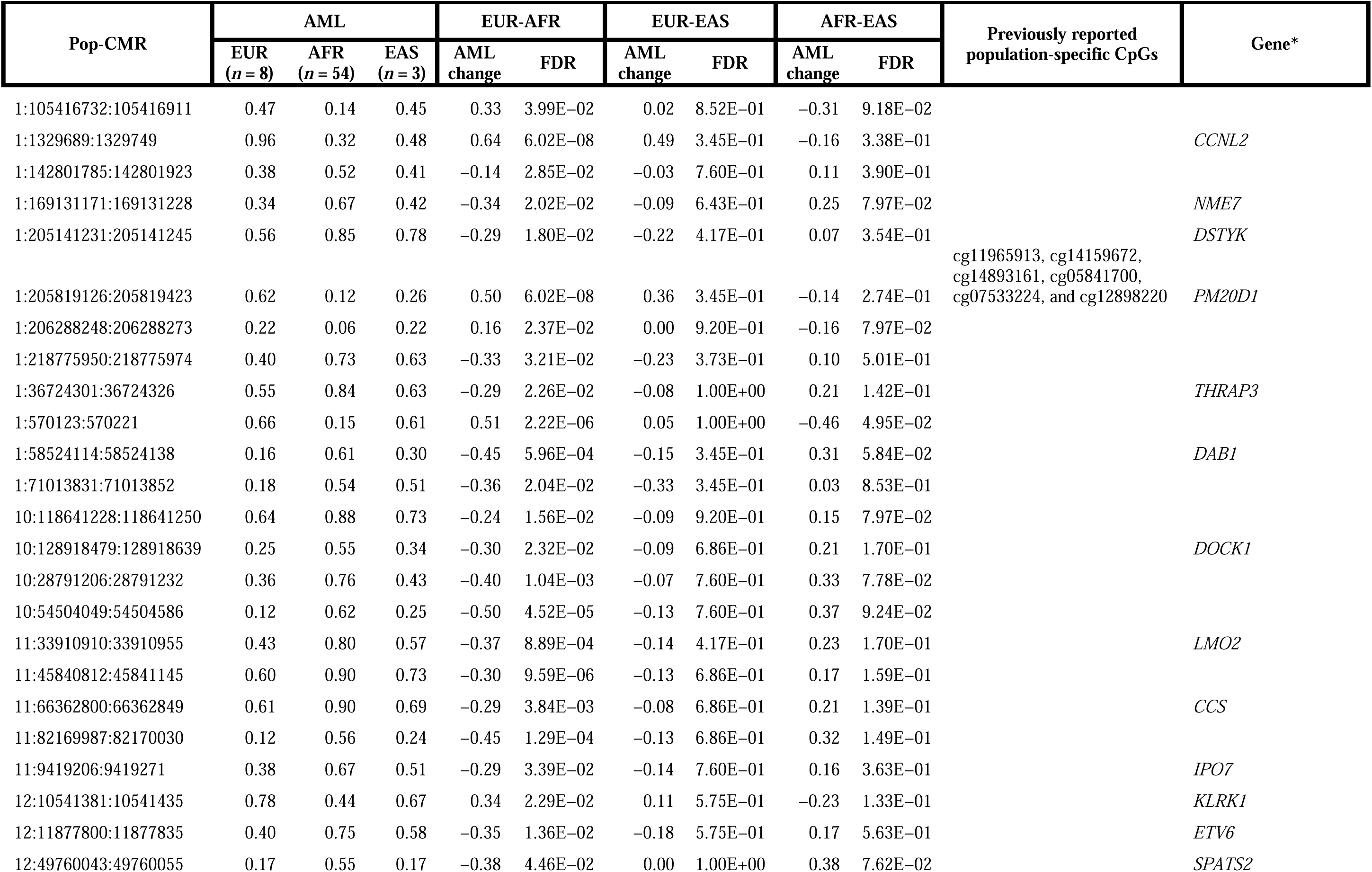
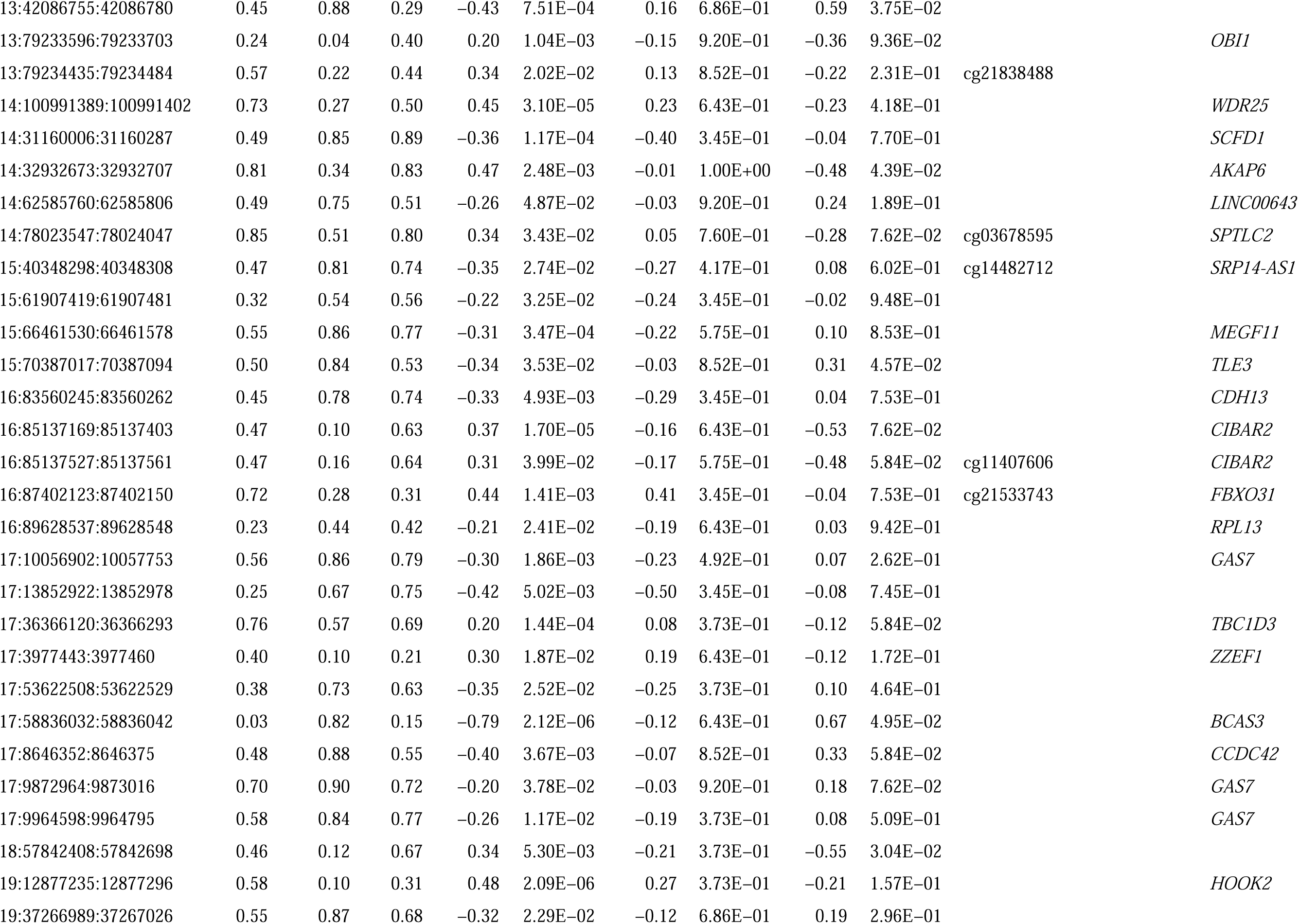
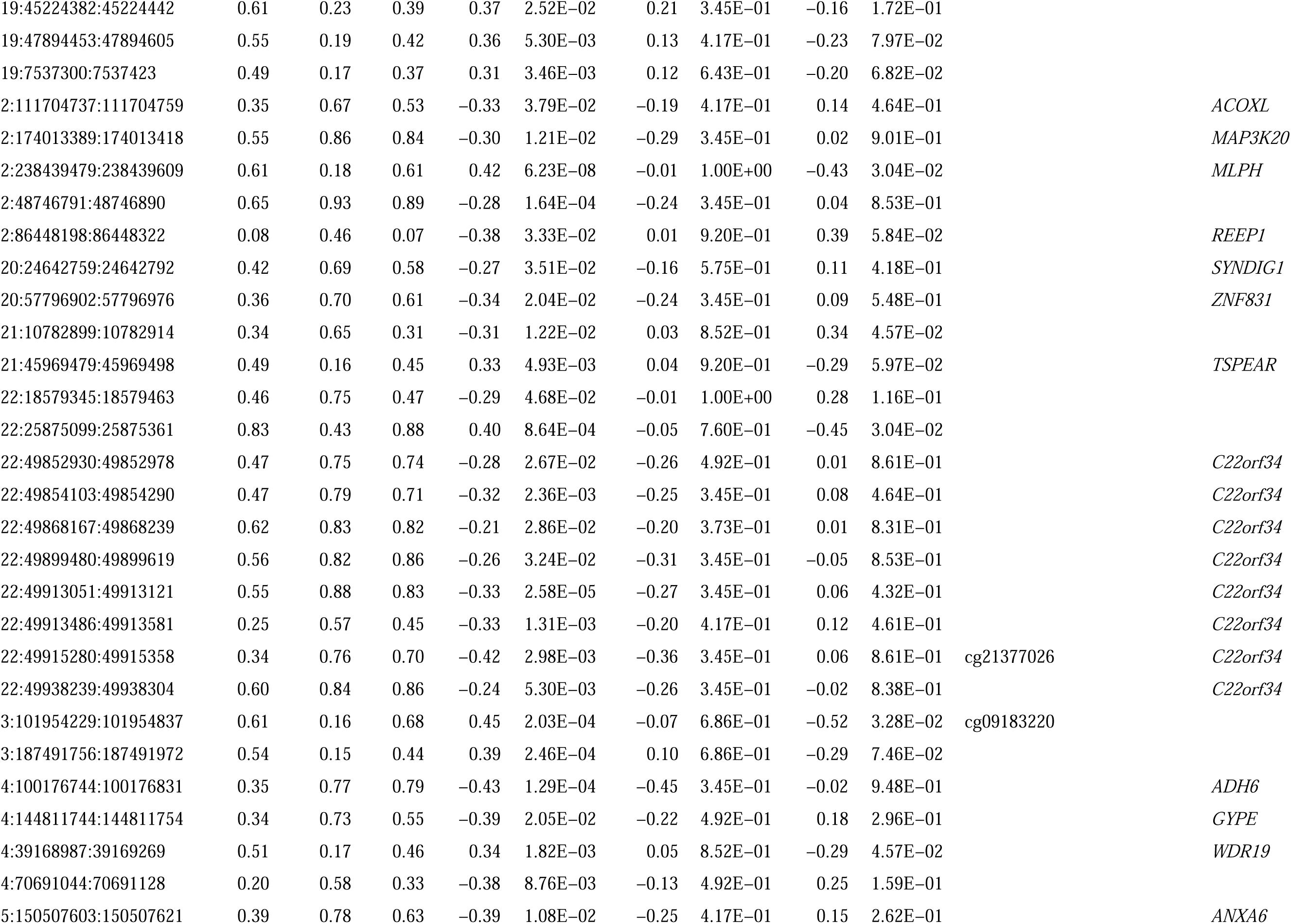
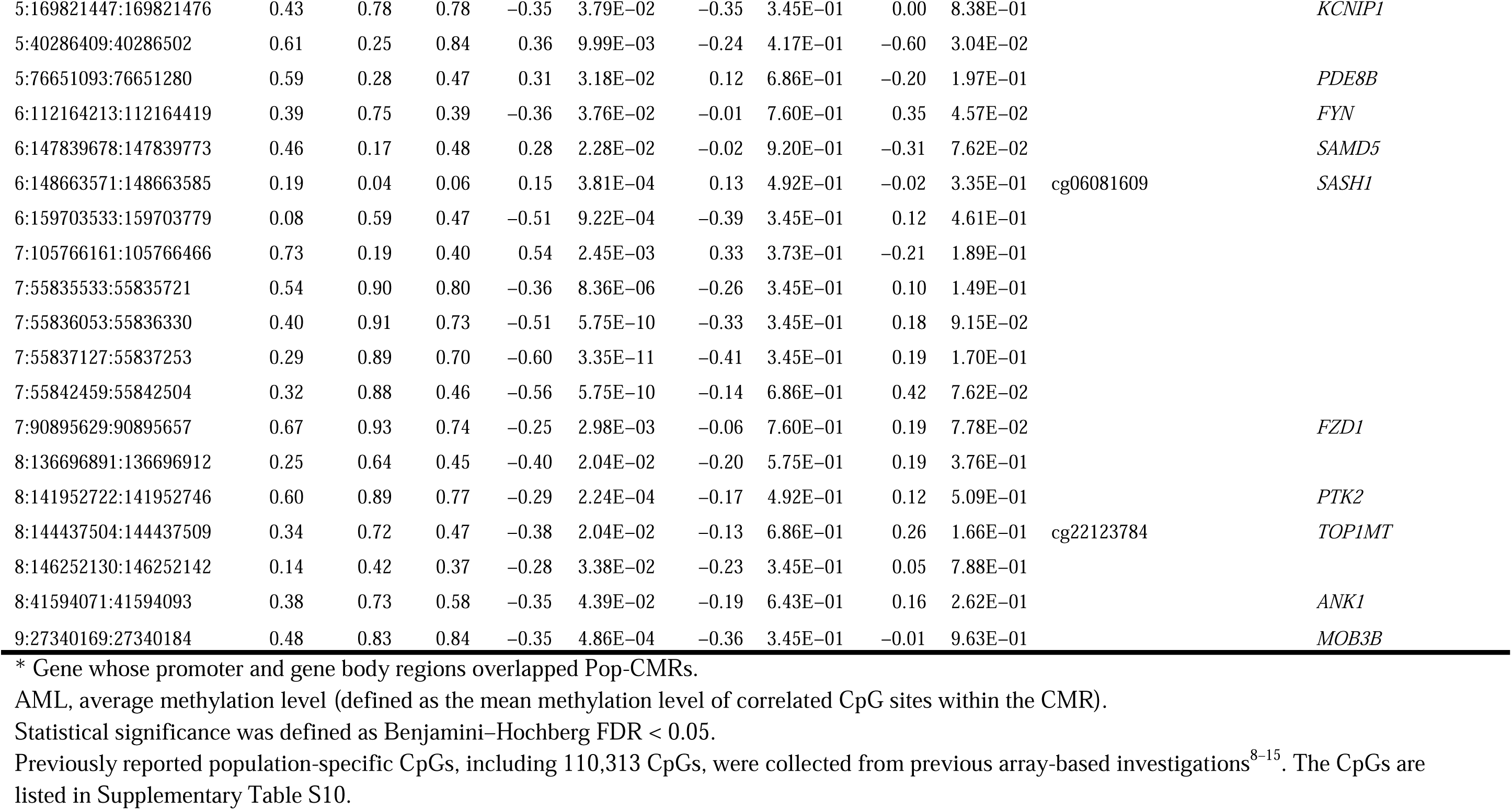
Methylation Changes of Population-Specific CMRs (Pop-CMRs) Across the Three Human Populations.

To further validate these Pop-CMRs, we performed an empirical study of Pop-CMRs in the combined HM450K data set described above containing 326 LCL samples (EUR *n* = 157, AFR *n* = 169) (data sources are described in Supplementary table S2) [12,14,30]. This analysis wanted to determine among all array CpGs (n = 413,012) whose DNAm levels were significantly different between populations, if there is an enrichment of these sites within Pop-CMRs (*n* = 7) compared with all sites (*n* = 413,012) or all sites within any CMRs (*n* = 3,936). Using a strict DNAm difference threshold of 0.1 and Mann–Whitney U test FDR < 0.05 [33], 57.1% (4/7) of CpGs in the array covered by Pop-CMRs showed statistically significant differences in DNAm between EUR and AFR populations with the same direction of change as observed in the WGBS data (Supplementary table S6), with a 17.4- and 111.0-fold increase in chance to be population-specific in comparison of random CpG sets from array CpGs covered by all CMRs and all CpGs in the array separately (1000 permutation test, both *P* < 0.001) (Fig. 2f). Notably, all seven CpGs examined here have been reported to show population-specific DNAm in the previous array-based investigations [9,16,34] (Supplementary table S7). These observations provided independent confirmation of a subset of Pop-CMRs with a larger sample size.

Additionally, we found that CpG count and density do not or very weakly influence the discovery of Pop-CMRs. The CpG count between Pop-CMRs and random CMR sets did not differ significantly (3.89 vs. 3.67, respectively, 1000 permutation test, *P* = 0.095). Moreover, there was no significant enrichment or depletion of Pop-CMRs in CpG-rich regions (e.g., CpG island) or CpG-poor regions (e.g., intron) compared to random sets of CMRs (1000 permutation test, all FDR > 0.05; Supplementary table S8), even though a nominally significant enrichment in introns and depletion in CpG islands were observed (both *P* < 0.05; Supplementary fig. S3). Interestingly, this result aligns with the findings obtained using the EPIC array [9] but contrasts with those using the HM450K array (i.e., population-specific DNAm sites were significantly depleted in CpG island) [12]. This suggested that leveraging a broader CpG coverage by, for example, using higher coverage microarray or WGBS, could yield more precise insights into the biological characteristics of population-specific DNAm.

#### Majority of population-specific DNAm patterns were exclusively found in our WGBS-based analysis compared to previous array-based studies

Although previous studies have identified population-specific DNAm sites using arrays, primarily the HM450K, one expected advantage of this study is the discovery of additional population-specific sites due to the comparatively high number of CpGs measured using WGBS. To highlight this significance, we examined the overlap and discrepancies of Pop-CMRs with previous array-based findings, including the HM27K, HM450K, and EPIC arrays. A total of 110,510 population-specific CpGs, mainly between EUR and AFR but also between other populations, were collected from previous array-based investigations regardless of tissue type [9–17] (the population-specific CpGs are listed in Supplementary table S9). Only 10 Pop-CMRs were found to overlap with these population-specific CpGs (Table 1). That is, DNAm patterns in 91 Pop-CMRs were uniquely discovered to be associated with population differences in this study, suggesting the major benefit of our whole-genome DNAm analysis in detecting DNAm alterations even with a limited sample size.

Comparing Pop-CMRs to all CpGs that can be measured in all arrays (HM27K, HM450K, and EPIC), only 15/101 were quantifiable (i.e., Pop-CMRs covering at least one CpG in the array) by at least one array (Fig. 2g). Five Pop-CMRs that overlapped with array CpGs had not previously been recognized as population-specific. Three of these were excluded due to technical artifacts of array probes, including cross-reactivity and the influence of SNPs on target CpG (i.e., probe adjacent to a SNP) [35–37] (Fig. 2g and Supplementary table S10). In WGBS data, sequencing reads that were aligned with a unique hit to the human reference genome were used to measure DNAm for these CpG sites due to the longer read length (length > 100 bp) than array probe sequence length (length = 50 bp). Unlike array DNAm measurements, which are modified by adjacent SNPs as their polymorphisms influence the mappability of the array probe [36,37], sequencing reads in WGBS were generated by chemistry cycles. The two remaining Pop-CMRs that overlapped with array CpGs, but were not previously identified as population-specific, may not have been detected due to the limited number of studies using the EPIC array. Only one study examined CpG DNAm changes between EUR and AFR individuals [9]. However, their multiple adjacent CpGs (distance < 15 kb) have been identified as population-specific CpGs in previous array-based investigations (Supplementary table S11). These results indicated that our whole-genome DNAm analysis offered additional benefits over previous array-based analyses, such as avoiding the influence of cross-reactivity and SNP polymorphism on array measurements, in detecting population-specific DNAm alterations.

Comparing genes associated with Pop-CMRs to those of population-specific EWAS findings is of particular interest as multiple CpGs in potentially different methylated regions could regulate the same genes. As such, the overlap between the genes containing population-specific CpGs discovered from previous array-based analyses was examined and compared to the 54 genes that overlapped with Pop-CMRs. Four genes (*CCDC42*, *GYPE*, *MAP3K20*, and *OBI1*) uniquely overlapped with Pop-CMRs alone, with three of these (*GYPE*, *MAP3K20*, and *OBI1*) being uniquely identified due to the absence of array probes measuring sites within these genes. These four genes have been reported to be associated with various diseases with differences in prevalence and incidence in different populations, such as type 2 diabetes mellitus [38], malaria infection [39], Alzheimer’s disease [40], and large artery stroke [41], suggesting the possible biological roles of the genes exclusively overlapping Pop-CMRs. Taken together, our investigation has led to the unique identification of a set of population-specific DNAm and gene methylation patterns linked to several metabolic and infectious diseases.

#### Pop-CMRs were enriched in genes relevant to metabolism and infection

Given the known role of DNAm in transcriptional regulation [9,42], we next performed pathway enrichment analysis for genes corresponding to Pop-CMRs to determine their possible biological relevance. Due to the limited amount of matched RNA sequencing data available, corresponding genes were identified in two ways: i.e., genes whose promoters or gene bodies overlapped Pop-CMRs; and genes whose transcription was significantly associated with the variation in DNAm of any CpGs comprising a Pop-CMRs using the iMETHYL database, which collected the results of WGBS and whole-transcriptome sequencing data for human peripheral blood mononuclear cells and blood cell types from Japanese subjects [43].

By overlapping Pop-CMRs to promoters or gene body regions, 54 unique genes were linked to 64 Pop-CMRs (Table 1). Pathway enrichment analysis showed significant enrichment of these genes in 318 biological processes with an FDR threshold of 0.05 and containing at least five tested genes (Supplementary table S12). These Gene Ontology (GO) pathways were classified based on ancestor GO terms [44], where the top three specific terms were metabolism process, biosynthesis process, and binding (Fig. 2h). In addition, consistent with the evolution of the B-cell immune response [45], we observed enrichments for certain pathways related to infection, such as response to stimulus (GO:0050896) and defense response (GO:0006952) (Supplementary table S12).

Using the iMETHYL database, the expression levels of 48 genes were found to be associated with 30 Pop-CMRs (Supplementary table S13), with 10 of these genes also identified in the GO analysis above (Fig. 2h). Genes whose expression levels were related to Pop-CMRs were significantly enriched in 214 biological processes with the same thresholds outlined above (Supplementary table S14). The top three ancestor GO terms of these biological processes were metabolism process, biosynthesis process, and cellular component (Fig. 2h). Additional enrichments included pathways related to immune function including response to stimulus (GO:0050896), immune response (GO:0006955), regulation of immune system process (GO:0002682), defense response (GO:0006952), leukocyte activation (GO:0045321), and infectious disease (REAC:R-HSA-5663205) (Supplementary table S14). The similar enrichment results for the two gene sets suggested the possible roles of population-specific DNAm in these specific biological processes, especially in metabolism and biosynthesis.

#### Increased between-population DNAm differences in Pop-CMRs linked to nearby genetic variants

DNAm levels at more than half of the population-specific CpGs discovered in this study (55.4%–70.2%) have been previously associated with DNA sequence variants [9–11,46]. However, these previous studies typically focused on CpGs measured in the arrays. As the majority of Pop-CMRs identified herein are not measured on any array, the extent to which the population specificity of CMRs was associated with genetic variation is largely unclear. To fill this gap, WGS genotype data sets corresponding to the 54 AFR WGBS samples were retrieved from the 1000 Genomes Project Phase 3 data set (data sources are described in Supplementary table S1) [47]. Associations between Pop-CMR AML and SNPs *in cis* (defined as SNPs located within a ± 10-kb window of each Pop-CMR [12,48]) were examined through linear modeling with adjustment for sex (Fig. 3a).

**Figure 3.**
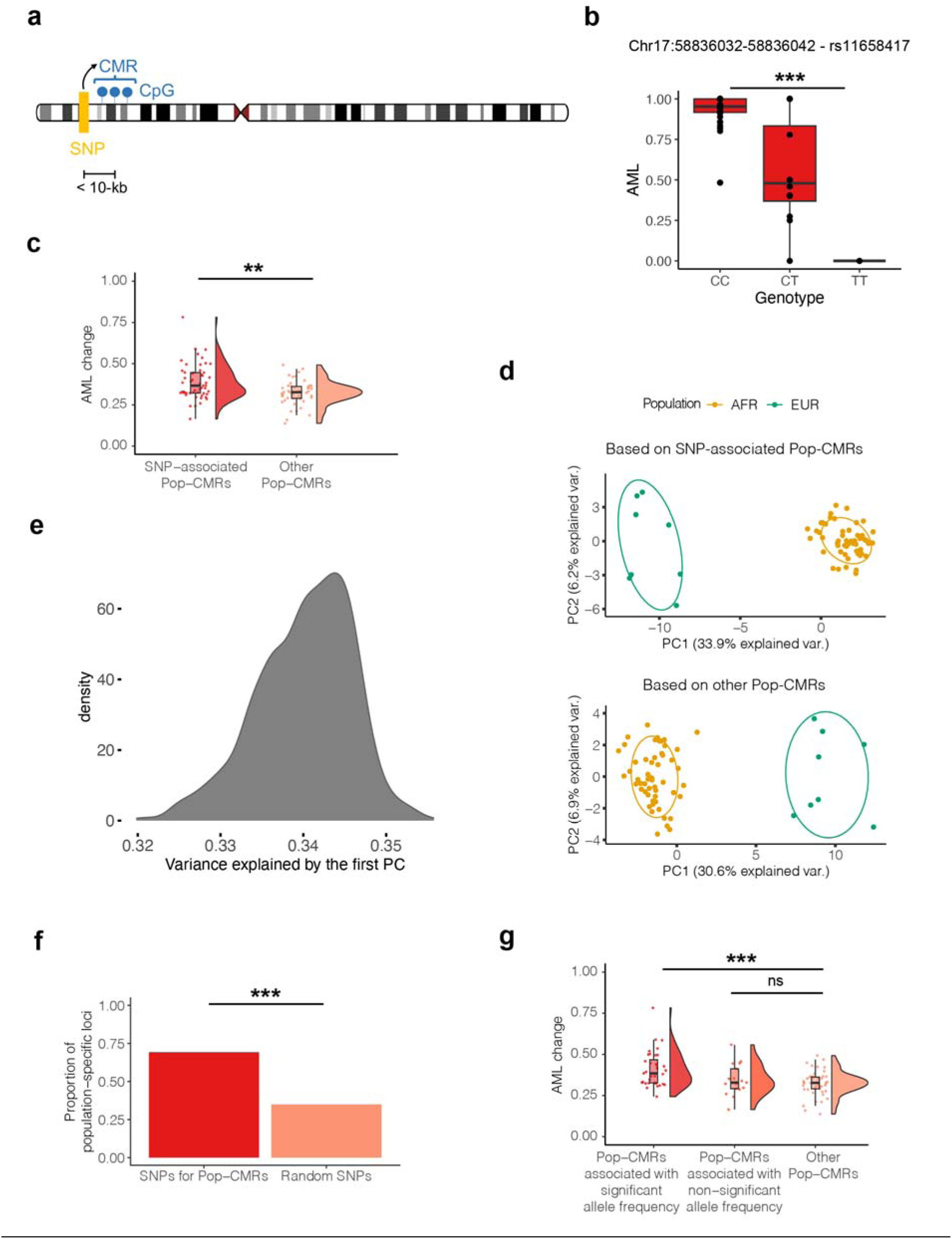
Interplay of Pop-CMRs with genetic variation. **(a)** Schematic of possible local genetic effects on CMR methylation (SNP-CMR distance < 10-kb (Heyn et al. 2013; Teh et al. 2014)). **(b)** Box plots of one SNP, rs11658417, showing significant association with methylation of one Pop-CMR (chr17:58836032–58836042). *** FDR < 0.001. **(c)** Box plots summarizing increased AML variability between EUR and AFR populations in SNP-associated Pop-CMRs. ** 0.001 ≤ *P* < 0.01. The absolute values of AML differences were used in the comparison and are illustrated in the figure. **(d)** Loadings using the first two principal components (PCs) of 53 SNP-associated Pop-CMRs and 48 other Pop-CMRs (independent of SNPs) for each WGBS LCL sample on principal component analysis (PCA) color-coded by ancestry. **(e)** The density of proportion of variance was explained by the first PC from AMLs of randomly selected SNP-independent Pop-CMRs. The random sampling process was repeated 1000 times. **(f)** Proportion of population-specific loci in SNPs for Pop-CMRs compared to random sets of SNPs. Random sampling was repeated 1000 times. *** *P* < 0.001. Population-specific loci were defined as SNPs with significant differences in allele frequencies between EUR and AFR populations based on Fisher’s exact test with an allele frequency difference threshold of 10% and an FDR threshold of 0.05. **(g)** Box plots summarizing AML variability between EUR and AFR populations in subgroups of SNP-associated Pop-CMRs in comparison of SNP-independent Pop-CMRs. *** FDR < 0.001, ^ns^ nonsignificant.

We found that 53 of 101 Pop-CMRs (52.5%) were associated with 707 SNPs (FDR < 0.05), consistent with results observed in DNAm array-based studies [9–11] (Supplementary table S15). Figure 3b shows an example of the DNAm patterns of the top-ranked Pop-CMR (chr17:58836032–58836042) with the largest AML change between EUR and AFR varying with the SNP, rs11658417. Given that multiple SNPs can be associated with the same CMR, only the most significantly associated SNP (i.e., with the lowest *P*-value) was used in subsequent analyses of each Pop-CMR (Table 2). These SNPs accounted for an average of 30.6% (range 14.7%– 64.4%) of the Pop-CMR DNAm variance, which was smaller than that reported in a previous array-based study in monocytes (58%) [9] but slightly larger than that in a previous array-based study in LCLs (26%) [11]. The findings suggested that the association between DNAm and population specificity may be influenced by sequence variants, particularly considering only the cis effects of SNP polymorphisms in this study.

**Table 2.**
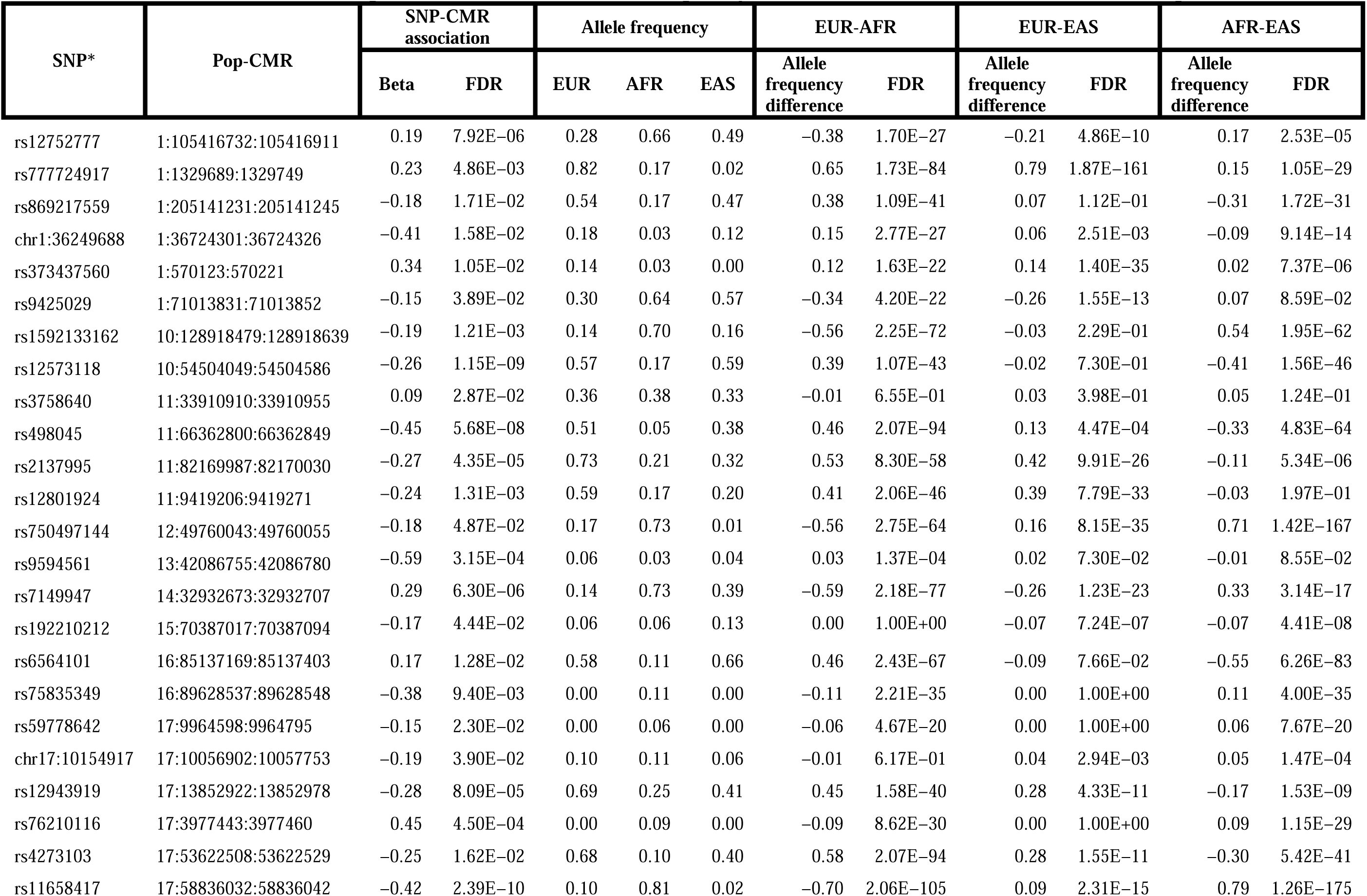
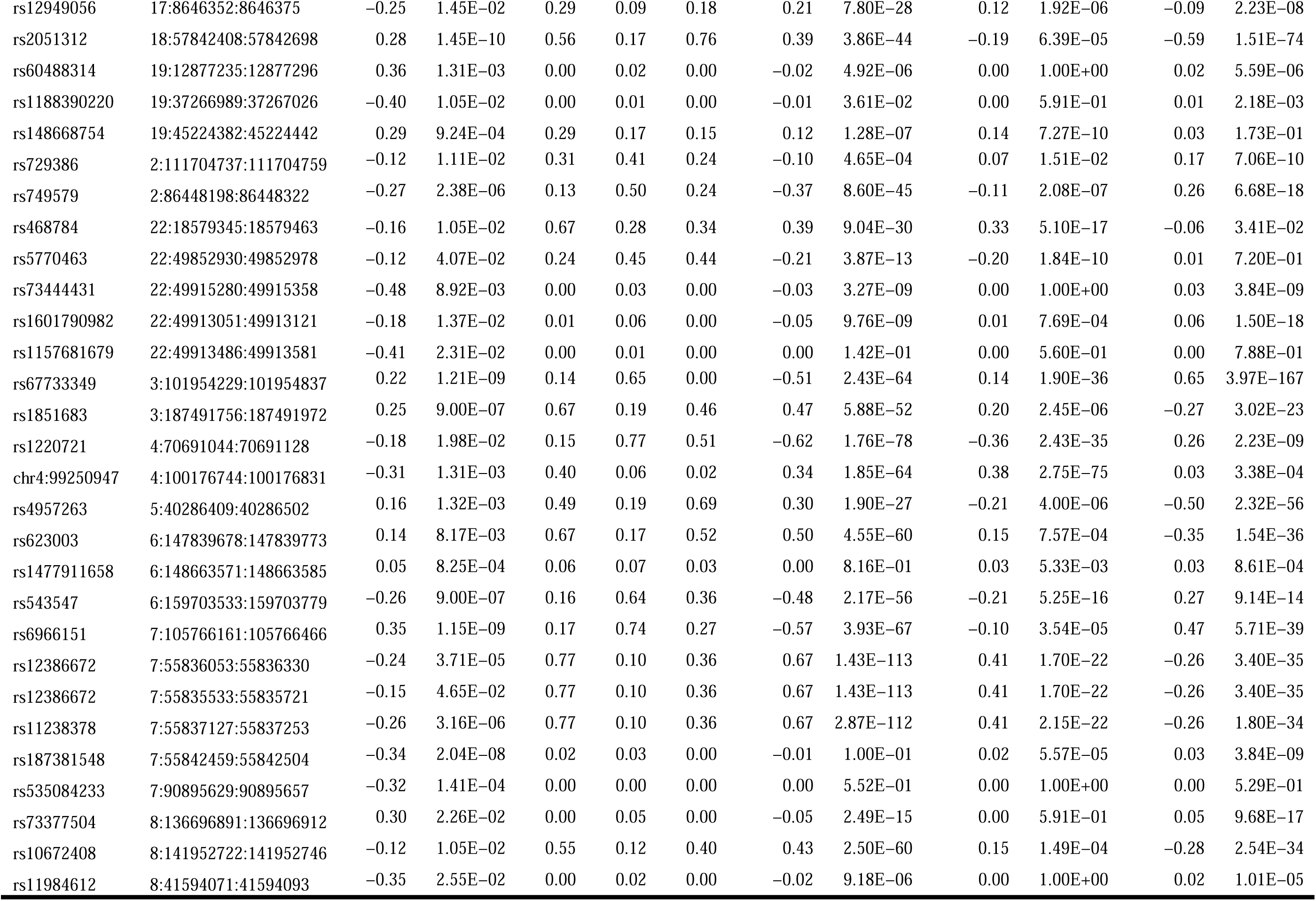
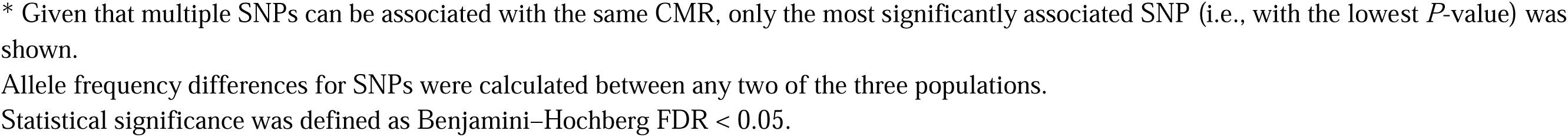
SNPs Associated With Pop-CMRs and Their Allele Frequency Differences Across the Three Human Populations.

It is important to study how genetic variants influence the population-specificity of DNAm, as an increase in DNAm differences between populations influenced by genetic variation may indicate a significant genetic contribution. We showed that the between-population DNAm differences were significantly larger for SNP-associated Pop-CMRs (*n* = 53) than other Pop- CMRs (*n* = 48) that were independent of SNPs (means: 0.39 vs. 0.32, respectively, Mann– Whitney U test *P* = 0.002) (Fig. 3c). Furthermore, PCA using the SNP-associated Pop-CMRs clearly distinguished the two populations, with the first PC accounting for 33.9% of the total variance (Fig. 3d). Although the SNP-independent Pop-CMRs could also clearly distinguish the two populations, they explained slightly less variance (30.6%). To address the concern that the difference in PC-explained variance may have been due to the unbalanced number of two sets of SNPs, a 1000 permutation test with SNP-independent Pop-CMRs and the same number of randomly selected SNP-associated Pop-CMRs was conducted. Similarly, the first PC for SNP-independent Pop-CMRs exhibited a significantly larger explained variance than that for SNP-independent Pop-CMRs (34.0% [range 32.0%–35.6%] vs. 30.6%, respectively, *P* < 0.001) (Fig. 3e). As these SNP-independent Pop-CMRs defined here may be associated with SNPs *in trans* or other genetic variations, such as copy number variants, these observations may be more significant when taking more genetic impacts into account. Taken together, our results suggested that genetic variation was associated with larger between-population DNAm differences in Pop- CMR, and hinted at a critical role of genetic variation for population-specific DNAm.

#### Allele frequency differences can explain the role of genetic variants in population-specific DNAm

We next explored whether the genetic influence on population specificity of DNAm could be attributed to differences in allelic frequency. First, we tested whether the genetic variants associated with the population specificity of DNAm exhibited varied allelic frequencies between populations. To this end, we examined allelic frequency differences in SNPs for Pop-CMRs between EUR (*n* = 503) and AFR (*n* = 661) populations using the 1000 Genomes Project Phase 3 data set (see Materials and Methods section “Investigation of the population specificity of SNPs for Pop-CMRs”) [47]. Of the 52 SNPs identified here (of which one SNP was associated with two Pop-CMRs), 36 (69.2%) showed significant allelic frequency differences between EUR and AFR populations (allelic frequency differences > 10%, Fisher’s exact test FDR < 0.05) (Table 2). This represented a ∼2.0-fold increase over the expected number of SNPs if frequency differences occurred randomly in the genome, suggesting enrichment of Pop-CMR-associated SNPs at loci with population-specific allelic frequencies (69.2% vs. 35.0%, respectively, 1000 permutation test, *P* < 0.001) (Fig. 3f).

Further, to ensure that the genetic influence on the population specificity of DNAm was due to population-specific SNP allelic frequencies rather than the SNPs themselves, we categorized Pop-CMRs into two groups based on the allelic frequency differences of the corresponding SNPs between populations (i.e., Pop-CMRs associated with SNPs showing significant population-specific allelic frequencies and those associated with SNPs showing no significant differences in allelic frequency) and then compared them to SNP-independent Pop-CMRs, respectively. The results revealed that Pop-CMRs associated with SNPs with significant population-specific allelic frequencies exhibited significantly greater between-population DNAm differences than SNP-independent Pop-CMRs (0.41 vs. 0.32, respectively, Mann–Whitney U test, FDR = 3.4 × 10^−4^), whereas Pop-CMRs associated with nonsignificant allelic frequency did not (0.35 vs. 0.32, respectively, Mann–Whitney U test, FDR = 0.589) (Fig. 3g). This demonstrated that the role of genetic variants in population-specific DNAm may be largely attributable to allelic frequency differences.

Notably, population-specific association (e.g., unique association in AFR but not EUR) between Pop-CMRs and SNPs may be another model indicating a role of genetic variants in population-specific DNAm. Due to the lack of genotype data sets corresponding to the EUR WGBS samples, we did not examine this model in the present study. Altogether, our findings suggested the potential substantial role of genetic variation in the population specificity of DNAm.

#### SNPs associated with Pop-CMRs were linked to metabolism and infection

Understanding the link between DNAm-associated genetic variants, referred to as methylation quantitative trait loci (mQTLs), and human phenotypes can point to disease-relevant pathways and contexts [9,49]. Here we evaluated the potential impacts of 707 SNPs associated with Pop- CMRs on complex traits by overlapping these SNPs with GWAS hits downloaded from the GWAS Catalog (last data release on 2023-01-30) [50]. The GWAS hits were filtered with a commonly used genome-wide significance threshold of *P* < 5.0 × 10^−8^, and all SNPs in high linkage disequilibrium (*r*^2^ = 0.8) with each of these hits were extracted based on the 1000 Genomes Project Phase 3 data set [47].

We found that 20 SNPs associated with six Pop-CMRs overlapped with GWAS hits related to various traits, such as obesity and cardioembolic stroke (Supplementary table S16). For instance, the SNP rs538656 (associated with Pop-CMR, chr18:57842408–57842698), which shows a notable difference in allele frequencies between EUR and AFR (allele frequency difference = 17.1% and Fisher’s exact test FDR = 5.9 × 10^−10^), has been reported to be linked to obesity susceptibility [51]. Based on 1000 permutations test, these SNPs exhibited stronger enrichment in GWAS traits, such as obesity and cardioembolic stroke, compared to random SNPs (Supplementary table S17).

Additionally, we conducted a pathway analysis for 32 genes whose promoters and gene bodies overlapped with SNPs associated with Pop-CMRs (Supplementary table S18). With an FDR threshold of 0.05, these genes were significantly enriched in 173 biological processes, each containing at least five tested genes (Supplementary table S19). These GO pathways were classified to ancestor GO terms [44], with the top three being metabolism process, biosynthesis process, and nucleobase-containing compound metabolic process (Supplementary fig. S4). These findings were consistent with our above findings involving genes whose promoters and gene bodies overlapped with Pop-CMRs, as well as genes whose expression was associated with Pop- CMR methylation. Similar enrichment results were also observed with regard to infection-related pathways, such as cellular response to stimulus (GO:0051716) and regulation of response to stimulus (GO:0048583) (Supplementary table S19). The overlap in biological processes found across comparisons suggests possible relations between population-specific DNAm, genetic variation, and complex traits.

#### A subset of Pop-CMRs showed population specificity beyond the test populations

Whether the population specificity of Pop-CMRs could be expanded to the other human populations in the 1000 Genomes Project [47], such as the EAS population, was explored (Fig. 4a). For this purpose, three EAS LCL WGBS samples were retrieved from GSE186383 (data sources are described in Supplementary table S1) [24]. The three populations formed distinct clusters on PCA using the 101 Pop-CMRs (Fig. 4b). We found that the population clustering was not due to chance, as PCA using random sets of CMRs in 1000 replicates showed no clustering based on population structure. We further examined the DNAm differences in Pop-CMRs between EAS and EUR samples, as well as between EAS and AFR samples. Thirteen Pop-CMRs showed significant differences between EAS and AFR populations (AML difference > 0.1 and Mann–Whitney U test FDR < 0.05) (Table 1). Although there were no significant DNAm differences between EAS and EUR populations, likely due to the limited number of samples from both populations, 72, 41, and 15 Pop-CMRs had AML differences greater than 0.1, 0.2, and 0.3, respectively, suggesting that a subset of Pop-CMRs may show population specificity for all three major human populations (Table 1).

**Figure 4.**
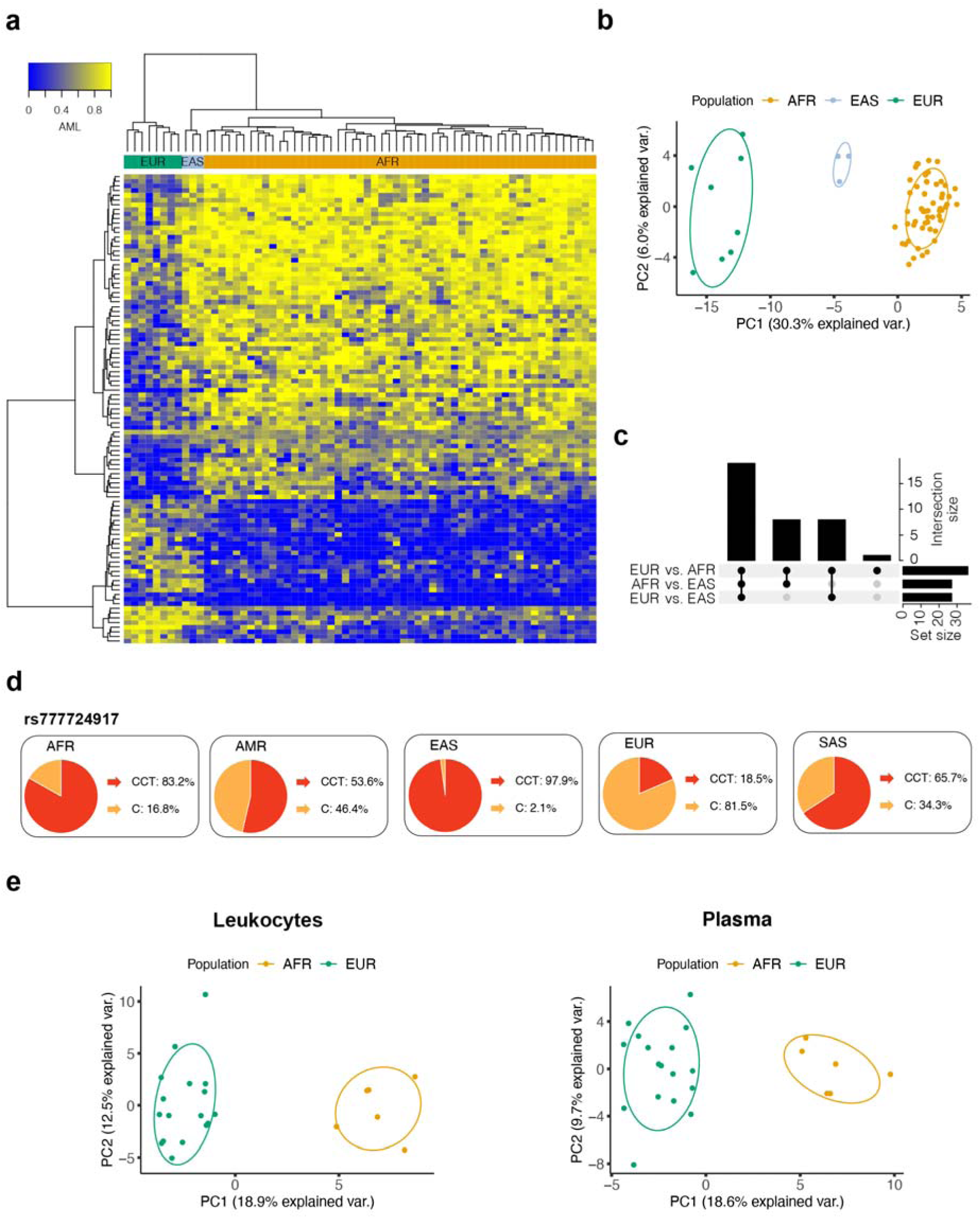
Expansion of 101 Pop-CMRs to the East Asian population and blood-based samples. **(a)** Unsupervised hierarchical clustering heat map using DNAm patterns from 101 Pop-CMRs. **(b)** Loadings using the first two principal components (PCs) of 101 Pop-CMRs for each WGBS LCL sample (AFR *n* = 54, EUR *n* = 8, EAS *n* = 3) on principal component analysis (PCA) color-coded by ancestry. **(c)** Upset diagram showing overlap of the differential allele frequency of SNPs for Pop-CMRs across three human populations. A total of 52 SNPs for Pop-CMRs were tested for allele frequency differences between EUR (*n* = 503), AFR (*n* = 661), and EAS (*n* = 504) populations in the 1000 Genomes Project. Differential allele frequencies were considered statistically significant between two populations at an allele frequency difference > 0.1 and FDR < 0.05. **(d)** The allele frequencies of rs777724917 in five human populations in the 1000 Genomes Project are shown in a pie chart. The reference allele is “CCT,” and the alternate allele is “C.” **(e)** Loadings using the first two PCs of 101 Pop-CMRs for each WGBS leukocyte sample (EUR *n* = 17, AFR *n* = 6) and each WGBS plasma sample (EUR *n* = 17, AFR *n* = 6) on PCA color-coded by ancestry.

As SNPs with allelic frequency differences between EUR and AFR populations play a substantial role in population specificity of Pop-CMRs, we speculated that the 52 SNPs associated with Pop-CMRs may also show differences in allelic frequency between EAS and EUR/AFR populations and thus present EAS-specific DNAm. With the genotype data from the 1000 Genomes Project (EUR *n* = 503, AFR *n* = 661, EAS *n* = 504) [47], we discovered that 27 and 27 SNPs associated with Pop-CMRs showed significant differences in allelic frequencies between EUR and EAS populations and between AFR and EAS populations, respectively (allele frequency difference > 10% and Fisher’s exact test FDR < 0.05) (Table 2). Of these, 19 SNPs exhibited significant differences in allele frequency between any two of the three populations and may underlie the observed DNAm differences between EAS and AFR populations (5 of 13 Pop-CMRs showing significant differences) or between EAS and EUR populations (7 of 15 Pop- CMRs showing DNAm differences greater than 0.3) (Fig. 4c). These results provided a genetic context for the potential value of Pop-CMRs in distinguishing these three human populations.

We further examined the allelic frequency differences of 52 SNPs associated with Pop-CMRs in two additional human populations from the 1000 Genomes Project: American (AMR *n* = 347) and South Asian (SAS *n* = 489) [52]. Using a threshold of allele frequency difference > 10% and Fisher’s exact test FDR < 0.05, we observed significant allele frequency differences for 9, 32, and 22 SNPs when comparing AMR with EUR, AFR, and EAS, respectively (Supplementary fig. S5). Additionally, 12, 33, and 23 SNPs showed differences when comparing SAS with EUR, AFR, and EAS, respectively. Notably, SNP rs777724917 (corresponding to Pop-CMR: chr1:1329689–1329749) displayed significant allelic frequency differences across all five populations (alternative allele [C] frequencies: 81.5% in EUR, 16.8% in AFR, 2.1% in EAS, 46.4% in AMR, and 34.3% in SAS) (Fig. 4d). Given the association of these SNPs with Pop- CMRs, it will be interesting to investigate whether Pop-CMRs show specific DNAm patterns in AMR and SAS populations.

#### Subsets of Pop-CMRs showed population specificity in peripheral blood leukocyte and plasma cell-free DNA samples

To assess the stability of Pop-CMRs in different tissues, peripheral samples were investigated as these are arguably more accessible and so would be more prevelant in future studies. The previously identified 101 Pop-CMRs were evaluated in peripheral blood leukocyte and plasma cell-free DNA samples from 23 healthy adult individuals (EUR *n* = 17, AFR *n* = 6) that were not included in the initial Pop-CMR identification process (data sources are described in Supplementary table S1) (GSE186888) [53]. In the blood leukocyte samples, PCA revealed that the two populations formed distinct clusters, with the first principal component, driven by these Pop-CMRs, accounting for 18.9% of the variance (Fig. 4e). We determined that this clustering was not due to chance, as PCA using random sets of CMRs in 1000 replicates showed no population clusters. Of the 101 Pop-CMRs, 22 were found to represent significant DNAm changes in leukocyte samples between the two populations (AML difference > 0.1 and Mann– Whitney U test FDR < 0.05) (Supplementary table S20).

Similar results were observed in plasma cell-free DNA samples. The two populations formed distinct clusters on PCA, with the first PC driven by these Pop-CMRs accounting for 18.6% of the variance (Fig. 4e). The population clustering driven by Pop-CMRs was not random compared to the results of PCA based on random sets of CMRs with 1000 replicates, which exhibited no clusters for population structures. Additionally, 21 of the 101 Pop-CMRs exhibited significant DNAm changes between the two populations in these plasma samples (AML difference > 0.1 and Mann–Whitney U test FDR < 0.05), with 17 of these also identified in leukocyte samples (Supplementary table S21). Taken together, these findings suggested that a subset of Pop-CMRs identified in LCLs may be extended to blood-based samples.

Among the 22 and 21 Pop-CMRs showing significant DNAm differences in leukocytes and plasma, 16 and 15 were associated with SNPs in LCL samples and showed differences in allelic frequencies between EUR and AFR populations, respectively (Supplementary tables S6, S16).

## Discussion

Comprehensively understanding genetic diversity, including epigenetics, is of fundamental interest in anthropological and medical sciences [2–5]. Initial studies of epigenetics have provided insights into the involvement of epigenetic variation in differences between human populations [9–16]. However, our understanding of epigenomic diversity between human populations is limited to the small proportion (∼3.0%) of the total CpGs in the genome that have been investigated to date. To address this limitation, DNAm differences between EUR and AFR populations using WGBS data were investigated. In addition, the co-methylation of adjacent CpGs was leveraged to improve the statistical power of the analyses, which was especially important given the comparatively small sample sizes investigated. Analysis identified 101 Pop- CMRs located in genes with potential functions related to metabolism and infection, where DNAm differences between populations were partly associated with underlying genetic variation. It is noteworthy that the Pop-CMRs identified between EUR and AFR LCL samples displayed excellent performance in not only population clustering of these samples but their clustering ability is expandable to the EAS population and peripheral blood-based tissues (i.e., leukocytes and plasma). The findings presented here will help us to obtain a better understanding of the epigenomic diversity between human populations, and thus facilitate the further development of tools for inferring population structures.

The major advantage of this study was the large number of CpGs measured by WGBS for the discovery of population-specific DNAm patterns. The results identified 86 Pop-CMRs that had not been evaluated previously in array-based investigations. More importantly, DNAm patterns in four genes (*CCDC42*, *GYPE*, *MAP3K20*, and *OBI1*) were identified exclusively here as being population-specific. These genes have been determined to be associated with various diseases with differences in prevalence and incidence in different populations, such as type 2 diabetes mellitus [38], susceptibility to malaria infection [40], Alzheimer’s disease [39], and large artery stroke [40]. These findings expand our knowledge of the role of epigenetics in population specificity and provide additional epigenetic insights to elucidate the susceptibilities and outcomes of certain diseases.

The importance of genetic variation in contributing to population-specific DNAm patterns was demonstrated in these results as more than half of Pop-CMRs were found to be associated with genetic variants in this study exhibiting compounded between-population differences in DNAm. This could be a consequence of genetic variation or genotype × environment interactions [10,11]. Here, a possible mechanism was suggested by which genetic variation may be driving population specificity of DNAm via differences in allele frequencies between populations, as noted in previous array-based studies and is now extended to WGBS-based studies [10,11,14]. More importantly, this mechanism may provide a genetic context to extend the population specificity of Pop-CMRs to other human populations and other tissue samples. As evidenced herein, population specificity of Pop-CMRs was successfully validated in the EAS population and two peripheral blood-based tissues, with empirical evidence that genetic variants can at least partly underlie these associations. Though genotype data for the peripheral-blood based tissue samples was unavailable making associations between Pop-CMRs and SNPs in these tissues impossible to validated, reassuringly, previous studies have indicated relatively low blood cell specificity of mQTLs [54,55]. As such, it is reasonable to speculate that the observed extension of Pop-CMRs to leukocytes and plasma samples may, at least in part, be due to the cross-tissue influence of population-specific genetic variants on DNAm. Although the findings regarding the extension of Pop-CMRs into new populations and tissues were not based on large data sets, the proposed genetic mechanism of population-specific DNAm lends confidence to these conclusions, can inform future studies, and potentially result in the advancing the development of tools for more accurate ancestry predictions based on this mechanism.

While the relations between Pop-CMRs and metabolic and infectious phenotypes were not explored directly in this study, multiple lines of indirect evidence suggested the possible association of Pop-CMRs with phenotypes related to metabolism and infection. Indeed, pathway enrichment analysis for genes mapped by Pop-CMRs, modified by Pop-CMRs, or mapped by Pop-CMR-associated SNPs all indicated considerable enrichment in metabolism- and infection-related processes. Furthermore, enrichment of SNPs for Pop-CMR with GWAS hits also showed the associations of these SNPs with obesity and cardioembolic stroke, the risk of which can be increased by infectious diseases [56]. Taken together, these observations suggest possible relations between Pop-CMRs and metabolic and infectious phenotypes. As immune responses have been shown to vary between human populations, these possible relations may be supported by the population-specific immune responses against infectious pathogens mediated by B cells, the functions of which are influenced by metabolic processes [2,57].

The WGBS data came from different laboratories increasing the risk for technical variation to be introduced. To address this, several approaches were applied to ensure any population-specific findings were robust. First, Non-Pop-CpGs (see Materials and Methods) were used to estimate and adjust for batch effects. Second, global CMR DNAm was determined to be very similar between EUR and AFR populations indicating minimal to no systematic technical artifacts, such as batch effects or variations in sample processing, affecting comparisons between sample populations. Third, Pop-CMRs which overlapped with array probes were mostly identified in previous studies, further suggesting the reliability of our approach (supplementary table S7) [9,16,34].

Although we identified CMR DNAm signatures unique to and discriminatory for EUR and AFR populations using the WGBS data sets and validated a subset of these signatures in DNAm array data with 326 samples (EUR *n* = 157, AFR *n* = 169), Pop-CMR identification was still limited due to the small sample size in this study. Furthermore, we assessed epigenetic changes across only three human populations, and extrapolation of the results to other populations will necessitate further analysis incorporating greater genetic diversity. Due to the small sample sizes, we were unable to accurately represent the individual or subpopulation diversity of each of the three human populations studied. It will be necessary to confirm whether these DNAm differences are robust to all individuals of EUR, AFR, and EAS ancestry. In addition, this study investigated and validated epigenetic differences only within LCLs and two peripheral blood-based tissues, even though it is well established that DNAm variation is considerably related to both tissue and cell type [58,59]. Further studies utilizing additional DNAm data sets with larger sample sizes, more diverse human populations, and across more tissues are required to obtain a more complete picture of pan-tissue DNAm pattern differences across human populations.

In summary, our analyses illustrated population-specific differences in whole-genome DNAm among individuals from three major human populations and elucidated their associations with genetic variation and biological relevance. This study provided crucial insight into epigenomic diversity across human populations and showcased its potential for inferring population structures.

## Materials and Methods

### WGBS data set information

A total of 70 WGBS LCL samples (EUR *n* = 9, AFR *n* = 58, EAS *n* = 3), 23 healthy leukocyte samples (EUR *n* = 17, AFR *n* = 6), and 23 healthy plasma samples (EUR *n* = 17, AFR *n* = 6) were retrieved from the BioProject database (PRJNA563344 and PRJNA733656) and the NCBI Gene Expression Omnibus (GEO) database (GSE186383, GSE57471, GSE66285, GSE89213, and GSE49627) for analysis (data sources are described in Supplementary table S1).). LCLs are produced by transforming primary B cells with Epstein–Barr virus (EBV) and are commonly used in immunology and human genetics research [60]. The details of sequencing were described in previous studies [23–28,53,61,62].

#### Genetic ancestry information confirmation

Genetic ancestry information was provided for all LCL samples, except for four individuals enrolled in a previous study from Karolinska Hospital (Stockholm, Sweden) [28]. To confirm the ancestry of these four subjects, we performed random forest prediction based on genotype information extracted from bisulfite sequencing reads by SNP calling in the WGBS data using BS-SNPer, a high-accuracy SNP caller based on a dynamic matrix algorithm and Bayesian statistical framework [20,63]. The parameters were set to call high-quality SNPs (options: -- minhetfreq 0.1 --minhomfreq 0.85 --minquali 15 --mincover 10 --maxcover 1000 --minread2 2 -- errorate 0.02 --mapvalue 10) where an average of 6,882,905 (range: 4,507,103–12,129,165) SNPs were found in each paired-end WGBS sample.

HapMap Phase 3 (release version 3) containing data for 1397 subjects from 11 ethnic groups (ftp://ftp.hgsc.bcm.tmc.edu/HapMap3-ENCODE/HapMap3/HapMap3v3) were randomly divided into training and validation data sets at a ratio of 4:1 [64]. Given that several populations have very similar genetic backgrounds, we combined the CHB (Han Chinese in Beijing, China), CHD (Chinese in Metropolitan Denver, CO, USA), and JPT (Japanese in Tokyo, Japan) into an EAS population and the CEU (Utah residents with Northern and Western European ancestry from the CEPH collection) and TSI (Tuscans in Italy) into a EUR population [64]. Furthermore, three individuals with known ancestry information (EUR *n* = 2, AFR *n* = 1) were included in the testing data set as positive controls. Prediction of ancestry was conducted using the `randomForest` R package, implementing Breiman’s random forest algorithm for classification [65]. Based on the accuracy of 0.853 for the prediction model with the validation data set, the ancestry of the four individuals with no reported ancestry information was predicted as EUR with 100% accuracy among positive controls.

#### WGBS data processing

WGBS data were preprocessed according to the Quality Control, Trimming, and Alignment of Bisulfite-Seq Data Protocol [66]. First, quality control (QC) checks on the raw WGBS data were applied using FastQC version 0.11.7 (https://www.bioinformatics.babraham.ac.uk/projects/fastqc/) [67]. To remove sequencing adapters and low-quality base reads, sequencing reads in paired-end and single-end WGBS files were trimmed with Trim Galore version 0.4.5 (https://www.bioinformatics.babraham.ac.uk/projects/trim_galore/) [68]. These trimmed reads were then mapped to the human genome (GRCh37/hg19) with Bismark version 0.19.0 (https://www.bioinformatics.babraham.ac.uk/projects/bismark/) using the default parameters [69]. In addition, the deduplication tool in Bismark was used to remove duplications in paired-end and single-end WGBS files. Finally, non-CG methylation reads were removed using the Bismark filtration tool (options: -s for single-end file; and -p for paired-end file) before identifying the CMRs. Five LCL WGBS samples (EUR *n* = 1, AFR *n* = 4) were excluded due to extreme low genomic coverage or becoming outliers on PCA checks.

#### Identification of CMRs from LCL WGBS

CpGs were extracted from sequence-aligned BAM files with highly mappable regions in the genome generated using a minimum mapping depth cutoff of 4 at more than half of the samples in each population [70]. Then, possible batch effects were corrected using 135 CpGs that were the top 5% of those stable between individuals across samples in each population (investigated based on the standard deviation of beta-converted M values [71]) and the top 5% of those stable between any two populations (all Mann–Whitney U test FDR > 0.1) (the CpGs are listed in Supplementary table S22). The FDR was calculated using the Benjamini–Hochberg method [72]. These CpGs were identified in DNAm array data with 288 LCL samples (EUR *n* = 96, AFR *n* = 96, EAS *n* = 96) from GSE36369 [12]. Their non–population-specific characteristics were further validated in WGBS samples before batch effect correction (Supplementary fig. S6). We used these non–population-specific and interindividual stable CpGs to estimate and regress out the DNAm changes caused by potential batch effects among WGBS samples. In addition, 13,462 CpGs associated with EBV transformation [28] and 168,914 CpGs in sex chromosomes were excluded to reduce the potential impact of EBV transformation and sex chromosome dosage effects on DNAm. Ultimately, 14,000,013 CpGs were retained for the identification of CMRs.

CMRs were identified according to the methods described in our previous report [21]. The methylation correlation of neighboring CpGs within a distance < 400 bp were examined across LCL samples using the Pearson method. CpGs with *r* > 0.5 and Benjamini–Hochberg FDR < 0.05 were considered significantly correlated. To reduce the potential effects of the unbalanced sample sizes of the EUR and AFR samples, a 1000 permutation test with EUR samples and the same number of randomly selected AFR samples was performed to validate these correlated CpGs with the same Pearson’s *r* and Benjamini–Hochberg FDR thresholds as above. To reduce the potential for false positives, only regions containing at least three highly correlated CpG sites were defined as CMRs.

#### HM450K data sets

Three HM450K data sets were used to further validate CMRs. The first cohort contained LCL samples retrieved from the GEO data portal under the accession number GSE36369 [12]. This data set consisted of 96 healthy EUR (Caucasian American) and 96 healthy AFR (African American) subjects from the Human Variation panel (sample sets HD100AA, HD100CAU, HD100CHI; Coriell Cell Repositories) and was collected by the National Institute of General Medical Science (NIGMS) [12]. The second HM450K data set included 133 HapMap LCLs derived from 60 EUR (Utah residents with Northern and Western European ancestry) and 73 AFR (Yoruba in Ibadan, Nigeria) individuals (GEO: GSE39672) [14]. The third data set was downloaded from GSE40699 and included five LCL samples (EUR [Utah residents with Northern and Western European ancestry] *n* = 4, AFR [Yoruba in Ibadan, Nigeria] *n* = 1) [30]. Four duplicate samples were removed across all three data sets. The remaining 326 HM450K LCL samples (EUR *n* = 157, AFR *n* = 169) were used for further analysis (data sources are described in Supplementary table S2). It should be noted that eight of these (EUR *n* = 1, AFR *n* = 7) were also subjected to bisulfite sequencing and were included in our LCL WGBS data.

#### DNAm array data preprocessing and analysis

For the array data sets used in this study, the downloaded data were normalized using the beta-mixture quantile (BMIQ) normalization method [73]. For each data set, the 65 SNP control probes, probes within the sex chromosomes, probes with missing values in more than 5% of samples, and polymorphic CpG probes were removed [36]. After filtration, 413,012 probes shared across the three data sets were retained for the analysis. Batch effects among the three data sets were corrected using *`ComBat`* from the `sva` R package [74]. The Pearson method was used to validate the correlation structures of CMRs identified from WGBS data. Similar to the analysis of WGBS data, a threshold of *r* > 0.5 and Benjamini–Hochberg FDR < 0.05 were used to define a significant correlation between CpGs.

#### Detection and validation of Pop-CMRs

WGBS data from 8 EUR and 54 AFR LCL samples were used for genome-wide analysis of CMR DNAm. The DNAm patterns of CMRs were measured as the AML of CpGs within each CMR. Differential AMLs between populations were determined by linear regression with adjustment for sex. The FDR was calculated using the Benjamini–Hochberg method [72]. CMRs with AML difference > 0.10 and FDR < 0.05 were defined as Pop-CMRs [33]. To evaluate and reduce the possible effects of the unbalanced sample sizes of the EUR and AFR samples, the 1000 permutation test with EUR samples and the same number of randomly selected AFR samples was conducted to validate Pop-CMRs. As AFR individuals in this study were recruited from two different subpopulations—Yoruba in Nigeria (*n* = 8) and Gambian in Western Division – Mandinka (*n* = 46)—and EUR samples were collected from two different countries—the USA (*n* = 5) and Sweden (*n* = 3)—1000 permutation tests were also performed to examine the Pop- CMR DNAm changes between subpopulations. Thirty-five CMRs that passed the AML and FDR thresholds were removed due to the possible effects of the unbalanced sample size (*n* = 34) and subpopulations (*n* = 1). PCA was performed to obtain a comprehensive view of these Pop- CMR DNAm patterns across samples using the *`prcomp`* function in the `stats` R package [75]. Validation of Pop-CMRs in array samples was performed using the Mann–Whitney U test with the same thresholds as above.

#### Enrichment analysis of CMRs, Pop-CMRs, and Pop-CMR-associated SNPs for known genomic elements

To calculate enrichment statistics for CMRs present in a certain genomic element, the same number of genomic regions with the same CpG count were randomly selected. The random sampling process was repeated 1000 times for each genomic element with fold enrichment calculated as the ratio of observed over expected values. Statistical significance was estimated based on the empirical *P*-value. Enrichments were considered statistically significant at a Benjamini–Hochberg FDR threshold of 0.05. Annotation files of coding exons, introns, 1 kb downstream, 5′-UTR, and 3′-UTR regions were retrieved from the UCSC Genome Browser (https://genome.ucsc.edu). LncRNA and pseudogene annotations were collected from the GENCODE database (https://www.gencodegenes.org/releases/19.html) [31]. Regulatory regions predicted from histone modifications using the ChromHMM algorithm were curated from the Encode LCL data set (http://hgdownload.cse.ucsc.edu/goldenPath/hg19/encodeDCC/wgEncodeBroadHmm) [76]. All genomic coordinates in this study were based on the human reference genome, GRCh37/hg19.

Similarly, 1000 permutations were implemented separately for analysis of genomic element enrichment within Pop-CMRs and Pop-CMR-associated SNPs. We calculated the fold enrichment of Pop-CMRs and SNPs in each genomic feature in comparison to randomly selected CMRs and SNPs, respectively. Enrichments were considered statistically significant at a Benjamini–Hochberg FDR threshold of 0.05.

#### Pathway enrichment analysis of genes

We mapped genes of interest to biological pathways to investigate their functional significance. These pathways included Gene Ontology (GO) molecular functions, GO biological processes, GO cellular components, the Kyoto Encyclopedia of Genes and Genomes (KEGG), and Reactome. These analyses were performed using the g:Profiler toolset (https://biit.cs.ut.ee/gprofiler/gost) [77]. A Benjamini–Hochberg FDR threshold of 0.05 was used to define statistical significance. To illustrate the basic GO term hierarchies and relations between terms within the context of biology, the free web tool, CateGOrizer, was used to investigate GO term data sets in terms of the GO classes they represented [44]. The “GO_Slim2” classification of GO terms was used (ftp://ftp.geneontology.org/pub/go/GO_slims/goslim_generic.obo).

#### Genotype data preprocessing

WGS genotype data corresponding to 56 samples (EUR *n* = 2, AFR *n* = 54) from WGBS data were downloaded from the 1000 Genomes Project Phase 3 [52]. QC checks, including checking discordant sex information, elevated outlying heterozygosity rate or missing genotype rate, high relatedness, and distinct population stratification, were conducted using PLINK v1.90b6 [78]. Two EUR individuals with an elevated outlying heterozygosity rate or missing genotype rate were removed. SNPs with minor allele frequency (MAF) < 0.01, missing genotype rate > 10%, or Hardy–Weinberg equilibrium exact test *P* < 1.0 × 10^−6^ were excluded [78]. Finally, 17,070,886 SNPs for 54 AFR individuals were retained for further analysis.

#### Mapping of SNP-associated Pop-CMRs

Genetically associated Pop-CMRs were scanned by testing SNPs located within a ± 10-kb window of each Pop-CMR, in line with previous studies indicating that *cis*-mQTLs are commonly located close to their corresponding SNPs [12,48]. The interplay between Pop-CMRs and neighboring SNPs was computed by a linear additive model with adjustment for sex using the `MatrixEQTL` R package [79]. Associations were selected at a Benjamini–Hochberg FDR threshold of 0.05.

#### Investigation of the population specificity of SNPs for Pop-CMRs

To understand how SNPs affect population-specific DNAm levels, the population specificity of 33 SNPs for Pop-CMRs was assessed using unrelated samples (EUR *n* = 503, AFR *n* = 661, EAS *n* = 504, AMR *n* = 347, and SAS *n* = 489) from the 1000 Genomes Project Phase 3 data set [47]. Population-specific loci were defined as SNPs with significant differences in allele frequency between two tested populations based on Fisher’s exact test with an allele frequency difference threshold of 10% and Benjamini–Hochberg FDR threshold of 0.05.

For each random set of SNPs, 33 SNPs were randomly selected from all SNPs (*n* = 17,070,886) after quality control and assessed for population specificity using the same approach as described for SNPs for Pop-CMRs. Based on 1000 permutations, fold enrichment was computed as the ratio of observed population-specific loci in SNPs for Pop-CMRs over randomly selected SNPs from background SNPs. Enrichment was considered statistically significant at an empirical *P*-value threshold of 0.05.

### Analysis of the expansion of Pop-CMRs in the East Asian population and blood-based

#### samples

Three Han Chinese LCL WGBS samples were collected from GSE186383 [24]. WGBS sequencing data were downloaded and preprocessed with the method described above. The potential batch effect was estimated and regressed out based on non–population-specific and interindividual stable CpGs. DNAm differences between EAS and EUR populations, as well as between EAS and AFR populations were calculated with the Mann–Whitney U test. DNAm changes with AML difference > 0.1 and Benjamini–Hochberg FDR < 0.05 were defined as statistically significant [33].

Blood-based samples were collected from GSE186888: leukocytes and plasma [53]. As raw data were not provided, processed BIGWIG files were downloaded. This data set included peripheral blood leukocyte samples and peripheral blood plasma samples collected from the same 23 individuals (EUR *n* = 17, AFR *n* = 6). CpGs within each Pop-CMR were extracted, and the AML was calculated for each Pop-CMR in each sample. Linear regression with adjustments for sex and age was performed to test Pop-CMRs’ associations with population specificity in leukocytes and plasma, separately. Associations with AML difference > 0.1 and Benjamini– Hochberg FDR < 0.05 were defined as statistically significant.

## Abbreviations

AFR: African
AMR: American
AML: average methylation level
BMIQ: beta-mixture quantile
CEU: Utah residents with Northern and Western European ancestry from the CEPH collection
CHB: Han Chinese in Beijing, China
CHD: Chinese in Metropolitan Denver, CO, USA
CMR: co-methylated region
CoMeBack: Co-Methylation with genomic CpG Background
DMR: differentially methylated region
DNAm: DNA methylation
EAS: East Asian
EBV: Epstein–Barr virus
EPIC: Illumina HumanMethylationEPIC BeadChip
EUR: European
EWAS: epigenome-wide association study
FDR: false discovery rate
GEO: NCBI Gene Expression Omnibus
GO: Gene Ontology
GWAS: genome-wide association study
HM27K: HumanMethylation27 BeadChip
HM450K: HumanMethylation450 BeadChip
JPT: Japanese in Tokyo, Japan
KEGG: Kyoto Encyclopedia of Genes and Genomes
LCL: lymphoblastoid B cell line
LncRNA: long noncoding RNA
MAF: minor allele frequency
NIGMS: National Institute of General Medical Science
PC: principal component
PCA: principal component analysis
Pop-CMR: population-specific CMR
QC: quality control
SAS: South Asian
SNP: single nucleotide polymorphism
TSI: Tuscans in Italy
WGBS: whole-genome bisulfite sequencing
WGS: whole-genome sequencing

## Declarations

## Acknowledgements

We are grateful to Compute Canada’s National Systems for Scientific Computation for computational support. We are extremely grateful to our colleague, Alan Kerr, whose wonderful contribution to writing and style greatly improved the manuscript.

## Author contributions

Z.D. conceived, designed, and performed the computational analyses, analyzed the data, and interpreted the results. N.G., M.F., S.S., and K.K. contributed ideas and participated in evaluating the results and discussions. M.S.K. supervised the study and obtained funding. Z.D. wrote the manuscript, with input from all authors. All authors approved the final version of the manuscript.

## Data availability

The WGBS, methylation array, and genotype data used in this study are available from the NCBI Gene Expression Omnibus (GEO; http://www.ncbi.nlm.nih.gov/geo/), the International HapMap Project (https://www.genome.gov/10001688/international-hapmap-project), and the 1000 Genomes Project (https://www.internationalgenome.org/).

## Code availability

All scripts used to generate the figures and support the findings of this study are available from the corresponding author upon reasonable request.

## Ethics approval and consent to participate

Not applicable.

