## Supplementary Figures for "Distinct Co-Methylation Patterns in African and European Populations and Their Genetic Associations"

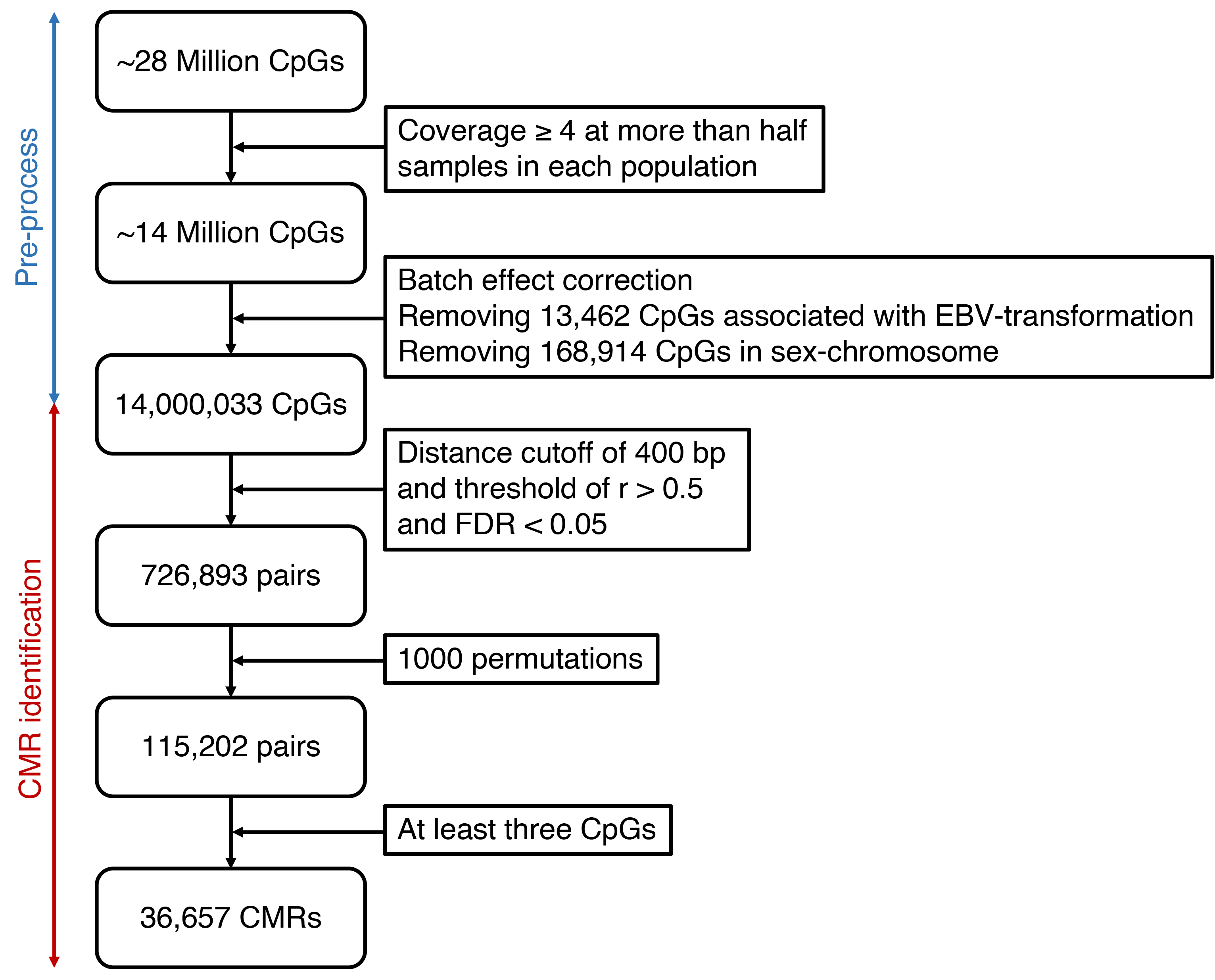

**Supplementary figure S1. CMR identification process pipeline.** A total of 62 WGBS samples (EUR *n* = 8, AFR *n* = 54) were used for the identification of CMRs after quality control. The alignment and depth cutoff processes were performed separately in each WGBS file. EBV, Epstein-Barr virus.

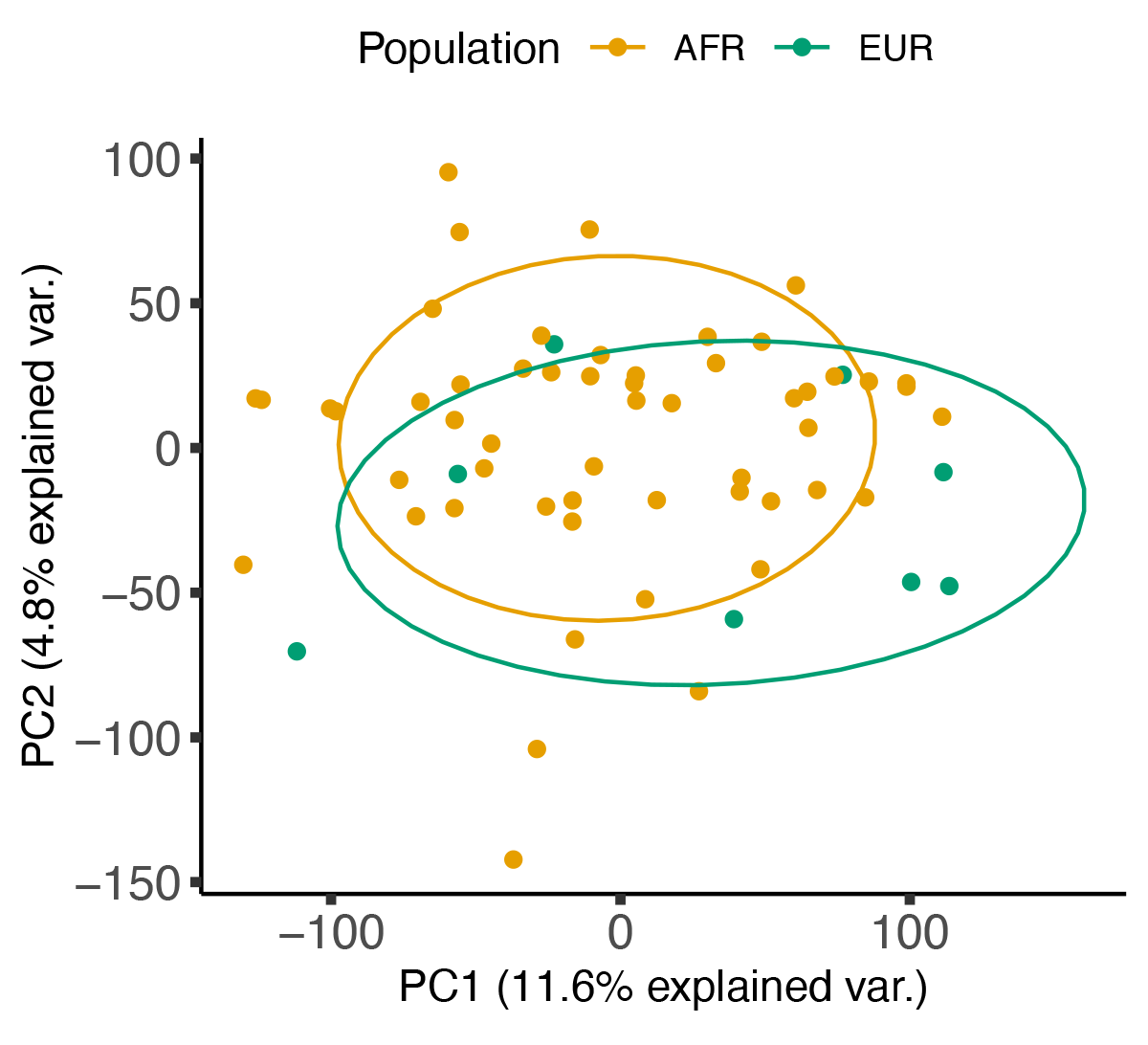

**Supplementary figure S2. Loadings using the first two PCs of all CMRs for each WGBS LCL sample (AFR *n* = 54, EUR *n* = 8) on principal component analysis (PCA) color-coded by ancestry.**

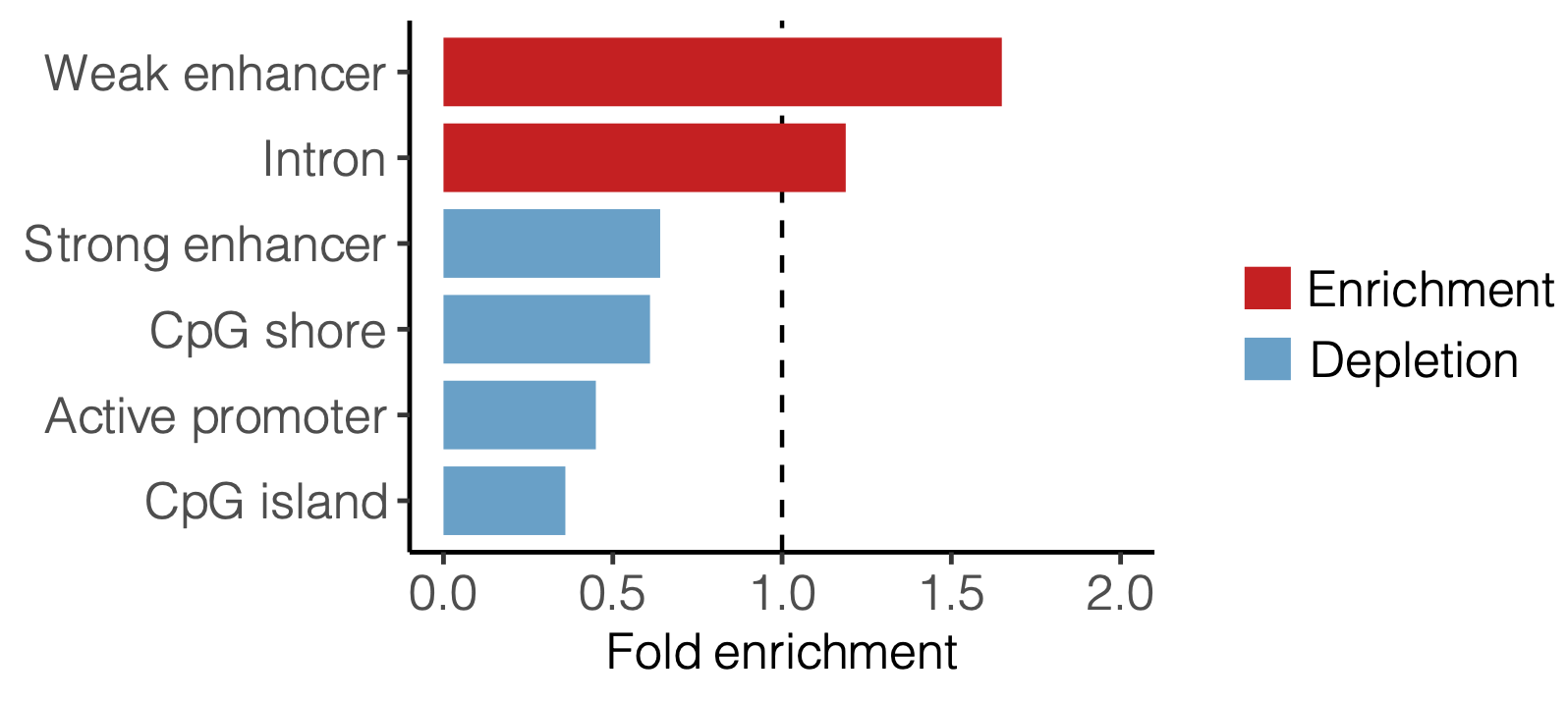

**Supplementary figure S3. Enrichment and depletion of 101 Pop-CMRs in genomic elements at nominal significance (*P* < 0.05).** Fold enrichment was calculated based on random sampling repeated 1000 times. For each set of CMRs enriched in a certain genomic element, the same number of CMRs were randomly selected.

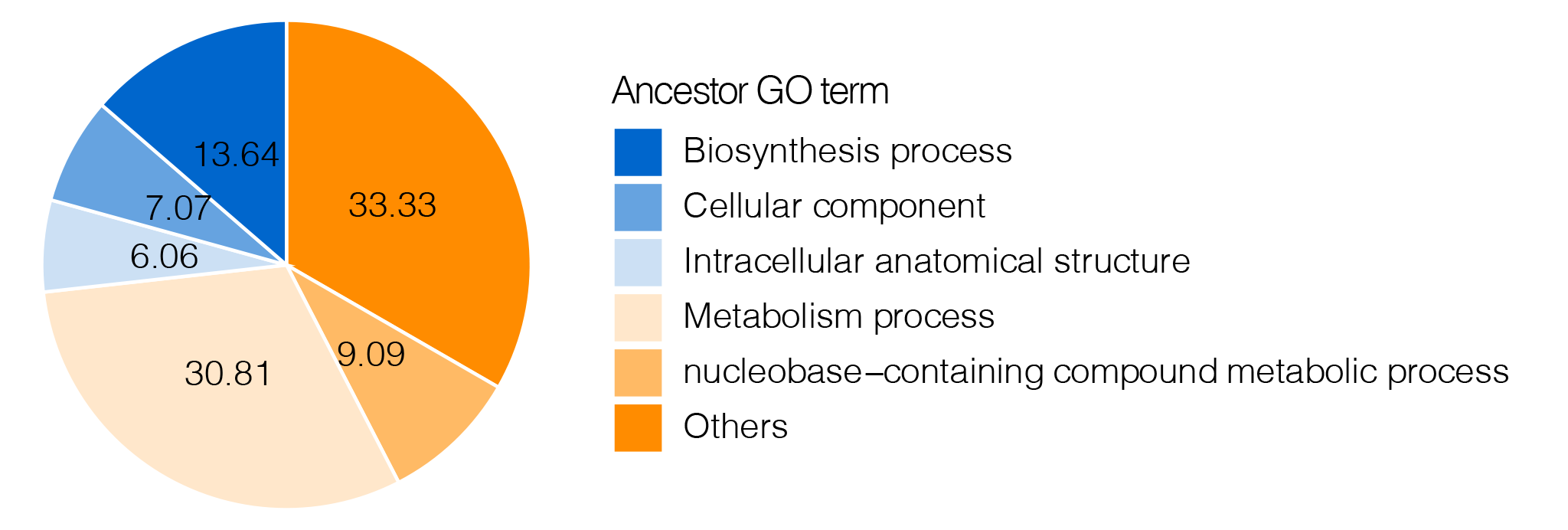

**Supplementary figure S4. Pie charts showing Gene Ontology (GO) classification for pathway enrichment analysis of 32 genes whose promoters and gene bodies overlapped SNPs associated with Pop-CMRs.**

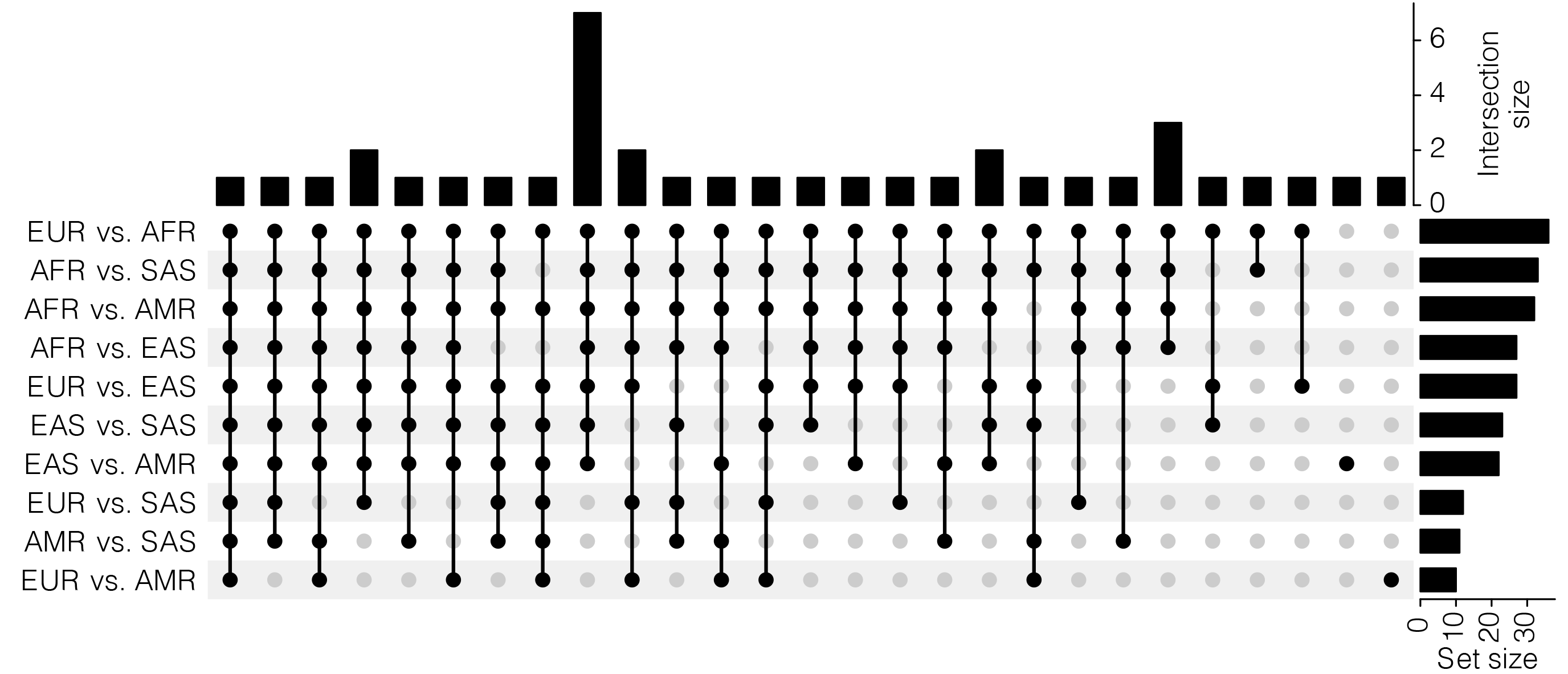

**Supplementary figure S5. Upset diagram showing overlap of differential allele frequency of SNPs for Pop-CMRs across three human populations.** A total of 52 SNPs for Pop-CMRs were tested for allele frequency differences between any two of five human populations (EUR *n* = 503, AFR *n* = 661, EAS *n* = 504, AMR *n* = 347, and SAS *n* = 489) in the 1000 Genomes Project. Differential allele frequencies were considered statistically significant between two populations at an allele frequency difference > 0.1 and FDR < 0.05.

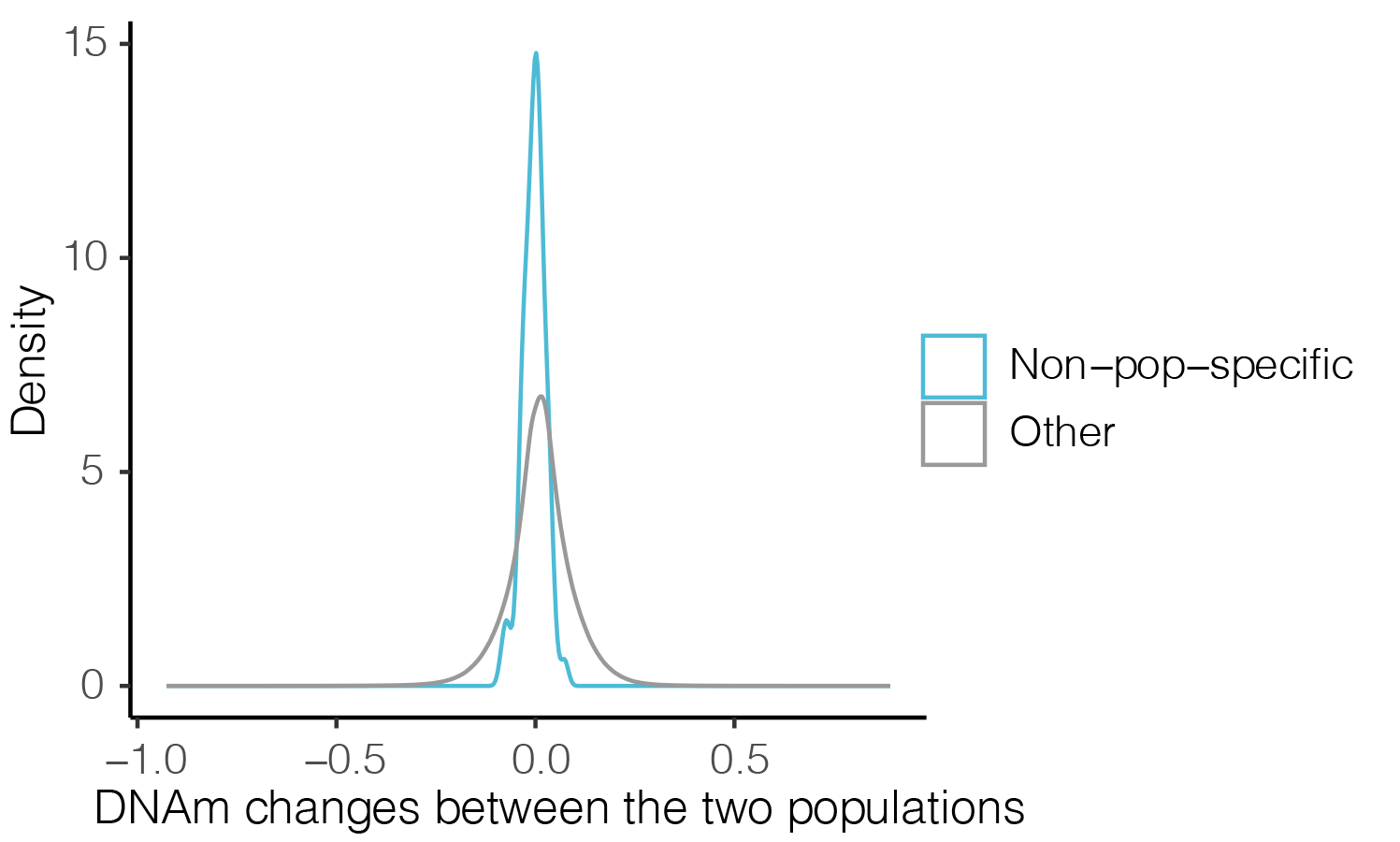

**Supplementary figure S6. Comparison of distributions of between-population DNAm changes among CpGs in WGBS data with and without population specificity.** These non–population-specific CpGs were identified in DNAm array data (EUR *n* = 96, AFR *n* = 96, EAS *n* = 96) as the top 5% of those stable between individuals across samples in each population and the top 5% of those stable between any two populations.
